# FBXO7/ntc and USP30 antagonistically set the ubiquitination threshold for basal mitophagy and provides a target for Pink1 phosphorylation *in vivo*

**DOI:** 10.1101/2022.10.10.511602

**Authors:** Alvaro Sanchez-Martinez, Aitor Martinez, Alexander J. Whitworth

## Abstract

Functional analyses of genes linked to heritable forms of Parkinson’s disease (PD) have revealed fundamental insights into the biological processes underpinning pathogenic mechanisms. Mutations in *PARK15/FBXO7* cause autosomal recessive PD and FBXO7 has been shown to regulate mitochondrial homeostasis. We investigated the extent to which FBXO7 and its *Drosophila* orthologue, ntc, share functional homology and explored its role in mitophagy *in vivo*. We show that *ntc* mutants partially phenocopy *Pink1* and *parkin* mutants and *ntc* overexpression supresses *parkin* phenotypes. Furthermore, ntc can modulate basal mitophagy in a Pink1- and parkin-independent manner by promoting the ubiquitination of mitochondrial proteins, a mechanism that is opposed by the deubiquitinase USP30. This basal ubiquitination serves as the substrate for Pink1-mediated phosphorylation which triggers stress-induced mitophagy. We propose that FBXO7/ntc works in equilibrium with USP30 to provide a checkpoint for mitochondrial quality control in basal conditions *in vivo* and presents a new avenue for therapeutic approaches.

## Introduction

Parkinson’s disease (PD) is the second most common neurodegenerative disorder. Autosomal recessive mutations in the genes encoding the mitochondrial kinase PINK1 (*PINK1*) and the E3 ubiquitin ligase Parkin (*PRKN*) are associated with early-onset parkinsonism. These genes have been shown to function in a common mitochondrial quality control pathway whereby mitochondria are degraded by autophagy (mitophagy). Briefly, PINK1 is partially imported to healthy polarised mitochondria where it is cleaved and subsequently degraded in the cytosol (1). Upon mitochondrial depolarisation, PINK1 is stabilised on the outer mitochondrial membrane (OMM) where it phosphorylates ubiquitin-Ser65 (pS65-Ub) conjugated to OMM proteins (2, 3). This acts as a signal for Parkin to be recruited and phosphorylated, relieving its autoinhibitory conformation, and allowing it to further ubiquitinate OMM proteins in close proximity (4, 5) that will serve as additional substrates for PINK1 creating a feed-forward mechanism (6). Counteracting mitochondrial ubiquitination, USP30, a mitochondrial outer membrane deubiquitinase, removes ubiquitin (Ub) from Parkin substrates acting as a negative regulator of mitophagy (7). The accumulation of pS65-Ub on the OMM triggers the recruitment of autophagy receptors, which promote autophagosome recruitment and, ultimately, degradation of the damaged mitochondria (8, 9).

However, most of our understanding of the function of PINK1 and Parkin come from the utilisation of mitochondrial uncouplers or inhibitors in cultured cells, usually in conjunction with Parkin overexpression (8, 10). Hence, there is a need to understand these mechanisms in more physiological model systems. *Drosophila melanogaster* models have delivered fundamental insights into the physiological function of the homologues *Pink1* and *parkin* (adopting FlyBase nomenclature), providing an excellent system to study mitochondrial homeostasis (11–16). Interestingly, while proteomic analysis of mitochondrial turnover in *Drosophila* supports a role of Pink1/parkin in mitochondrial quality control (17), studies using pH-sensitive fluorescent mitophagy reporters showed that Pink1 and parkin had minimal impact on basal mitophagy (18–21). Thus, many questions endure regarding the mechanisms of action of Pink1/parkin in mitochondrial quality control *in vivo* and the potential contribution of other key players.

Mutations in the gene encoding F-box protein 7 (*FBXO7)*, have been found to cause autosomal recessive early-onset parkinsonism similar to those caused by mutations in *PINK1* and *PRKN* (22, 23), motivating a need to understand how FBXO7 is related to PINK1/Parkin biology. F-box domain-containing proteins are essential factors of the SCF-type (Skp1-Cul1-F-box) E3-ubiquitin ligase complexes, which are responsible for recruiting Ub target substrates via the F-box domain (24). FBXO7 has both SCF-dependent and SCF-independent activities (25, 26). Although it has been previously shown to cooperate with PINK1 and Parkin in mitophagy (27), how this occurs *in vivo* is poorly characterised warranting further investigation in an animal model. *Drosophila* encode a single homologue of *FBXO7* called *nutcracker* (*ntc*), which was identified in a screen for genes that control caspase activation during a late stage of spermatogenesis (28). ntc shares sequence, structure, and some functional similarities with mammalian FBXO7 (26, 28, 29) and has been shown to regulate proteasome function via its binding partner PI31, an interaction that is conserved in FBXO7 (28, 29). Thus, *Drosophila* provides a genetically tractable system to study role of ntc/FBXO7 in mitochondrial quality control *in vivo*.

Mitochondrial quality control mechanisms are believed to have an important pathophysiological role in PD. However, how this process is orchestrated *in vivo* and the upstream factors involved are still unclear. Therefore, we sought to investigate the relationship between ntc, Pink1 and parkin, and their role in mitophagy, in particular basal mitophagy. We have found that ntc is able to compensate for the absence of *parkin* but not *Pink1*. Moreover, ntc plays an important role in promoting basal mitophagy in a Pink1- and parkin-independent manner, which is opposed by its counteracting partner USP30. This mechanism sets a threshold for basal mitophagy by regulating the steady-state levels of ubiquitin on OMM proteins, subsequently modulating the levels of pS65-Ub. Together, we provide evidence of a novel checkpoint for basal mitophagy that could potentially serve as therapeutic target for neurodegenerative disorders.

## Results

### Overexpression of *ntc* can rescue parkin but not Pink1 mutant phenotypes

The F-box domain localised in the C-terminal of *Drosophila* ntc shares considerable sequence similarity with that of human FBXO7 (28). Both proteins have a conserved valine residue involved in substrate recognition, including common interacting partners, e.g. PI31 (26, 29). However, a functional conservation in between FBXO7 and ntc has not been formally tested *in vivo.* Notably, it has been shown that FBXO7 functionally interacts with mammalian PINK1 and Parkin and genetically interacts with *parkin* in *Drosophila* (27). Therefore, as an initial exploration of the functional homology between ntc and FBXO7, we tested for a similar genetic interaction between *ntc* and *parkin. Drosophila parkin* mutants present multiple disease-relevant phenotypes, including locomotor deficits, dopaminergic neurodegeneration, and a severe mitochondrial disruption leading to flight muscle degeneration (13, 14). Strikingly, overexpression of *ntc* by the strong ubiquitous *daughterless* (*da*)-GAL4 driver significantly rescued these *parkin* phenotypes, (Figure 1A-1D). At the molecular level, we observed that ntc is also able to decrease the steady state levels of the *Drosophila* Mitofusin (MFN1/MFN2) homologue, Marf, which has been previously shown to be increased in *parkin* mutants (16) (Figure 1E). These results support a functional homology between FBXO7 and ntc, and, moreover, indicate that ntc can at least partially substitute for parkin *in vivo*.

**Figure 1.**
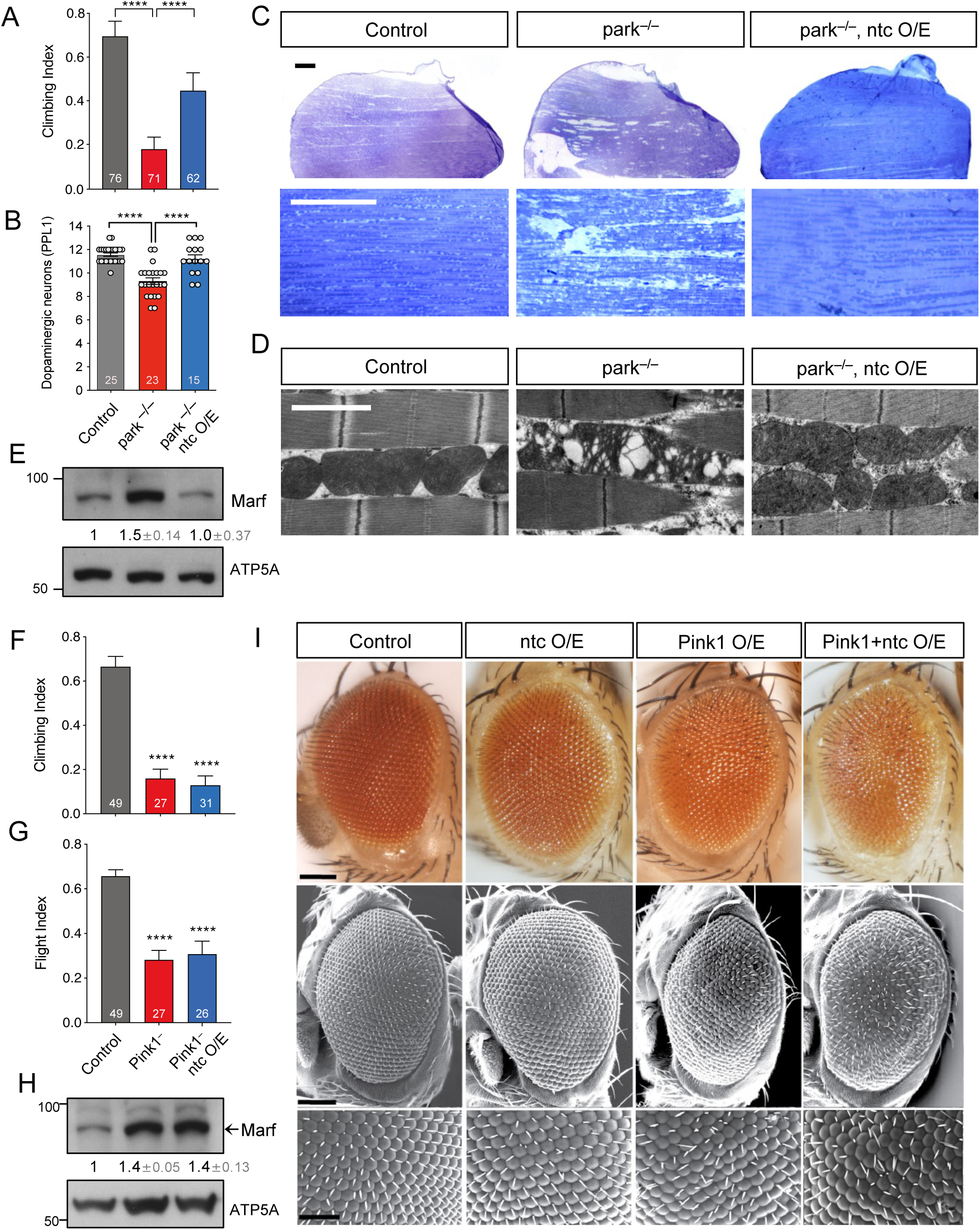
ntc overexpression is able to rescue *parkin* but not *Pink1* mutants. (**A**) Climbing ability of 2-day-old control and *parkin* mutants alone or with transgenic expression of *ntc* with the ubiquitous driver *da*-GAL4. Chart show mean ± 95% CI and n values. Kruskal-Wallis nonparametric test with Dunn’s post-hoc test correction for multiple comparisons; **** *P* < 0.0001. (**B**) Quantification of dopaminergic neurons (PPL1 cluster) in 30-day-old control and *parkin* mutants alone or with transgenic expression of *ntc* with the ubiquitous driver *da*-GAL4. Data represented as mean ± SEM; n shown in chart. One-way ANOVA with Bonferroni post-hoc test correction; **** *P* < 0.0001. (**C**) Toluidine blue staining, and (**D**) transmission electron microscopy (TEM) analysis of thoraces from 2-day-old control and *parkin* mutant alone or with transgenic expression of *ntc* with the ubiquitous driver *da*-GAL4. Scale bars; top = 200 μm, middle = 10 μm, bottom = 2 μm. (**E**) Immunoblot analysis from whole fly lysates of Marf steady state levels from 2-day-old control and *parkin* mutants alone or with transgenic expression of *ntc* with the ubiquitous driver *da*-GAL4. Numbers below blots indicated the mean ± SD of Marf levels normalised to ATP5A loading control across 3 replicate experiments. (**F**) Climbing and (**G**) flight ability of 2-day-old control and *Pink1* mutants alone or with transgenic expression of *ntc* with the ubiquitous driver *da*-GAL4. Chart show mean ± 95% CI; n shown in chart. Kruskal-Wallis nonparametric test with Dunn’s post-hoc test correction for multiple comparisons; **** *P* < 0.0001. (**H**) Immunoblot analysis from whole fly lysates of Marf steady state levels from 2-day-old control and *Pink1* mutants alone or with transgenic expression of *ntc* with the ubiquitous driver *da*-GAL4. Numbers below blots indicated the mean ± SD of Marf levels normalised to ATP5A loading control across 3 replicate experiments. (**I**) Analysis of the compound eye by (top panels) light microscopy or (middle and bottom panels) scanning electron microscopy (SEM) of 2-day-old control, overexpression of ntc, Pink1 or both combine using the *GMR*-GAL4 driver. Scale bars; top and middle = 100 μm, bottom = 50 μm.

In contrast, assessing the functional relationship between ntc and Pink1 in a similar manner, *ntc* overexpression was unable to rescue climbing or flight locomotor defects of *Pink1* mutants, nor the increased Marf steady-state levels (Figure 1F-1H). Furthermore, the neurodegenerative ‘rough eye’ phenotype – commonly used for testing genetic interactors (30, 31) – induced by *Pink1* overexpression, in a robustly stereotyped manner, was not enhanced by *ntc* overexpression as previously shown for *parkin* (15). These results mirror equivalent analyses of *FBXO7* (27), and indicate that *Pink1* and *ntc* do not obviously genetically interact.

### *ntc* mutants partially phenocopy *Pink1/parkin* mutants

If ntc performs similar functions to the Pink1/parkin pathway in *Drosophila*, we reasoned that *ntc* mutants may phenocopy *Pink1/parkin* mutants. Supporting this, *ntc* mutants, like *Pink1* and *parkin* mutants, are male sterile due to defective sperm individualization (11–13, 28, 32). To extend this, we assessed *ntc* mutants for classic *Pink1/parkin* phenotypes as described above. *ntc* mutants homozygous for an amorphic nonsense mutation (*ntc*^ms771^; abbreviated as *ntc*^−/−^) showed a marked defect in climbing and flight ability (Figure 2A, 2B). We verified this was likely specific to *ntc* by assessing a hemizygous condition (*ntc*^−/Df^) with comparable results (Figure 2A, 2B). Importantly, transgenic re-expression of *ntc* was able to significantly rescue the climbing and flight deficit (Figure 2A, 2B), confirming that the phenotype is specifically caused by loss of ntc function. Also similar to *Pink1/parkin* mutants, *ntc* mutants showed a dramatically shortened lifespan, with median survival of ~7 days compared to ~55 days for controls (Figure 2C), which was almost completely rescued by transgenic re-expression of *ntc* (Figure 2C).

**Figure 2.**
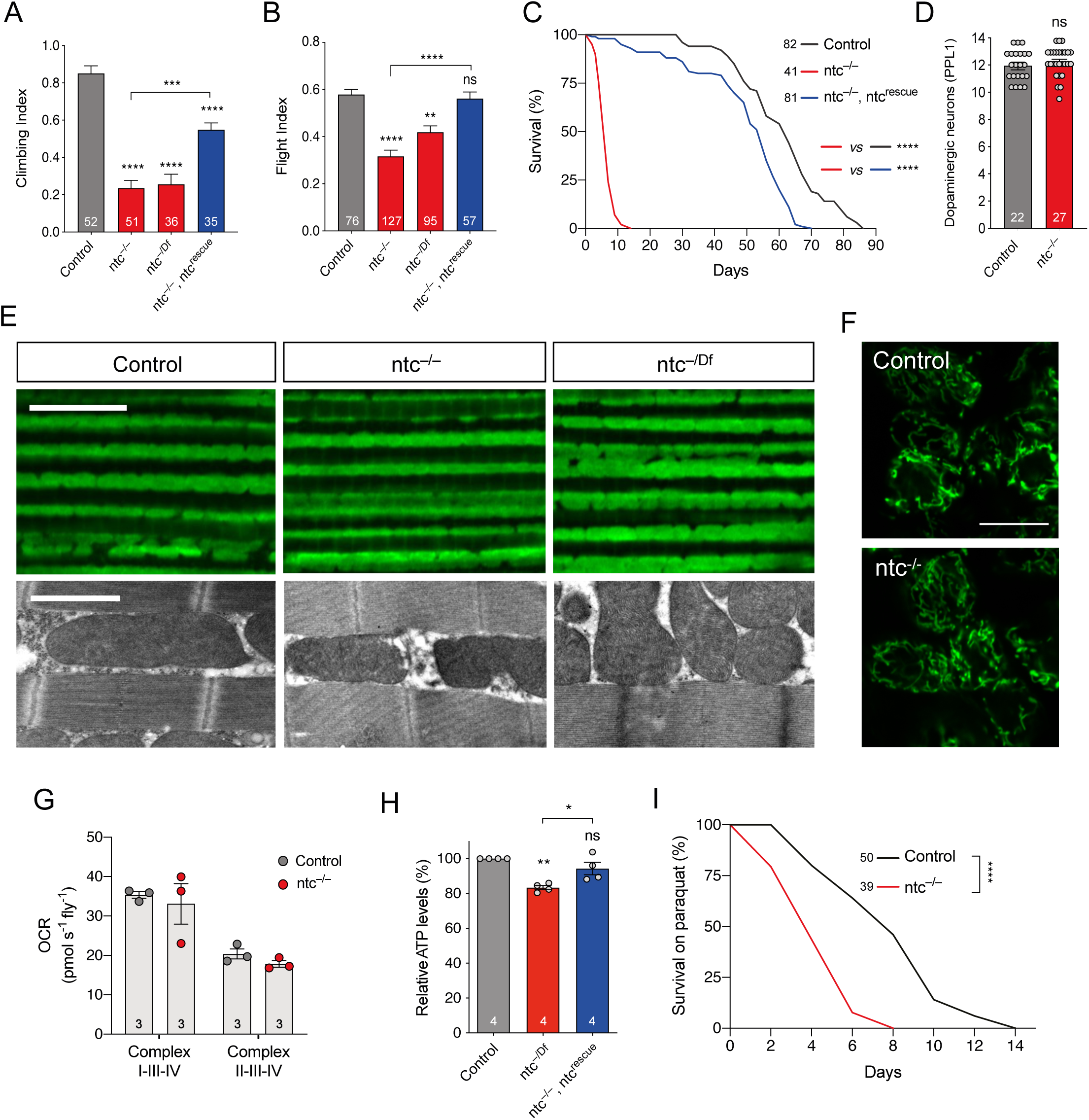
Loss of ntc has motor and lifespan deficits, no gross effect on mitochondria but increased sensitivity to oxidative stress. Analysis of (**A**) climbing and (**B**) flight abilities of 2-day-old control and *ntc* mutants alone or with transgenic expression of *ntc* with the ubiquitous driver *da*-GAL4. Chart show mean ± 95% CI and n values. Kruskal-Wallis nonparametric test with Dunn’s post-hoc test correction for multiple comparisons; *\*\* P* < 0.01, *\*\*\* P* < 0.001, **** *P* < 0.0001. (**C**) Lifespan assay of control and *ntc* mutants alone or with transgenic expression of *ntc* with the ubiquitous driver *da*-GAL4; n shown in chart. Log rank (Mantel-Cox) test; **** *P* < 0.0001. (**D**) Quantification of dopaminergic neurons (PPL1 cluster) in 10-day-old control and *ntc* mutants alone. Data represented as mean ± SEM; n shown in chart. One-way ANOVA with Bonferroni post-hoc test correction. (**E**) (Top panel) Immunohistochemistry with anti-ATP5A staining and (bottom panel) transmission electron microscopy of 2-day-old adult thoraces from control and *ntc* mutants. Scale bars; top = 10 μm, bottom = 2 μm. (**F**) Mitochondrial morphology in motoneurons from the ventral nerve cord of third instar larvae in control and *ntc* mutants, overexpressing mito-GFP with the pan-neuronal driver *nSyb*-GAL4. Scale bars = 10 μm. (**G**) Oxygen consumption rate from complex I and complex II of 2-day-old control and *ntc* mutants. Data represented as mean ± SEM; n shown in chart. One-way ANOVA with Bonferroni post-hoc test correction. (**H**) ATP levels of 2-day-old control and *ntc* mutants alone or with transgenic expression of *ntc* with the ubiquitous driver *da*-GAL4. Data represented as mean ± SEM; n shown in chart. One-way ANOVA with Bonferroni post-hoc test correction; * *P* < 0.05, *\*\* P* < 0.01. (**I**) Lifespan assay of control and *ntc* mutants treated with 10 μM paraquat. n shown in chart. Log rank (Mantel-Cox) test; **** *P* < 0.0001.

However, in contrast to *Pink1/parkin* mutants, we did not observed loss of DA neurons in the PPL1 clusters of *ntc* mutants, although the short lifespan limited the analysis to only 10-day-old flies (Figure 2D). Likewise, immunohistochemical analyses of *ntc* mutants revealed no apparent disruption of mitochondrial morphology or integrity at confocal or electron-microscopy levels in flight muscle (Figure 2E) or larval central nervous system (CNS) (Figure 2F). Analysing mitochondrial respiratory capacity, *ntc* mutants were not consistently different from controls despite a downward trend (Figure 2G). Nevertheless, ATP levels were reduced in *ntc* mutants (Figure 2H), which was rescued by re-expression of wild-type *ntc*.

Oxidative stress is a consistent feature of PD pathology and normal ageing. Many models of PD show sensitivity to oxidative stressors such as paraquat (PQ), including *Drosophila parkin* mutants (33, 34). Similarly, *ntc* mutants also showed increased sensitivity to PQ exposure (Figure 2I). Together, these results show that loss of *ntc* causes previously undescribed phenotypes that impact organismal vitality similar to, but also different from, *Pink1/parkin* mutants, though the effects are generally less severe.

### ntc regulates basal mitophagy and antagonises USP30 in a Pink1/parkin-independent manner

Previous work showed that FBXO7 facilitates PINK1/Parkin-mediated mitophagy upon mitochondrial depolarisation in cultured cells (27). Thus, we next sought to determine whether *Drosophila* ntc acts to promote mitophagy *in vivo*. The *mito*-QC reporter provides a sensitive and robust reporter for mitophagy whereby the GFP of OMM-localised tandem mCherry-GFP is quenched in the acidic lysosome, revealing mitolysosomes as ‘mCherry-only’ puncta (35). Importantly, in the absence of potent mitochondrial toxins – not readily applicable *in vivo* – the steady-state analysis of *mito*-QC provides an insight into basal mitophagy (20, 36). Quantifying basal mitophagy in larval neurons revealed a significant reduction in *ntc* mutants compared to controls, which was rescued upon re-expression of *ntc* (Figure 3A, 3B). Conversely, *ntc* overexpression significantly increased basal mitophagy (Figure 3C, 3D). To verify these results, we analysed an alternative mitophagy reporter with a mitochondrial matrix-targeted version of *mito-QC*, termed *mtx-QC*, with equivalent results (Figure S1A, S1B). Thus, ntc is both necessary and sufficient to promote basal mitophagy. Similarly, transgenic expression of *FBXO7* also increased mitophagy in larval neurons, underscoring their shared functionality (Figure S2A, S2B). Further validating these results in human cells stably expressing the *mito*-QC reporter (37), siRNA-mediated knockdown of *FBXO7* also significantly reduced basal mitophagy (Figure S2C-S2E). Together these results support a conserved function of FBXO7/ntc in regulating basal mitophagy.

**Figure 3.**
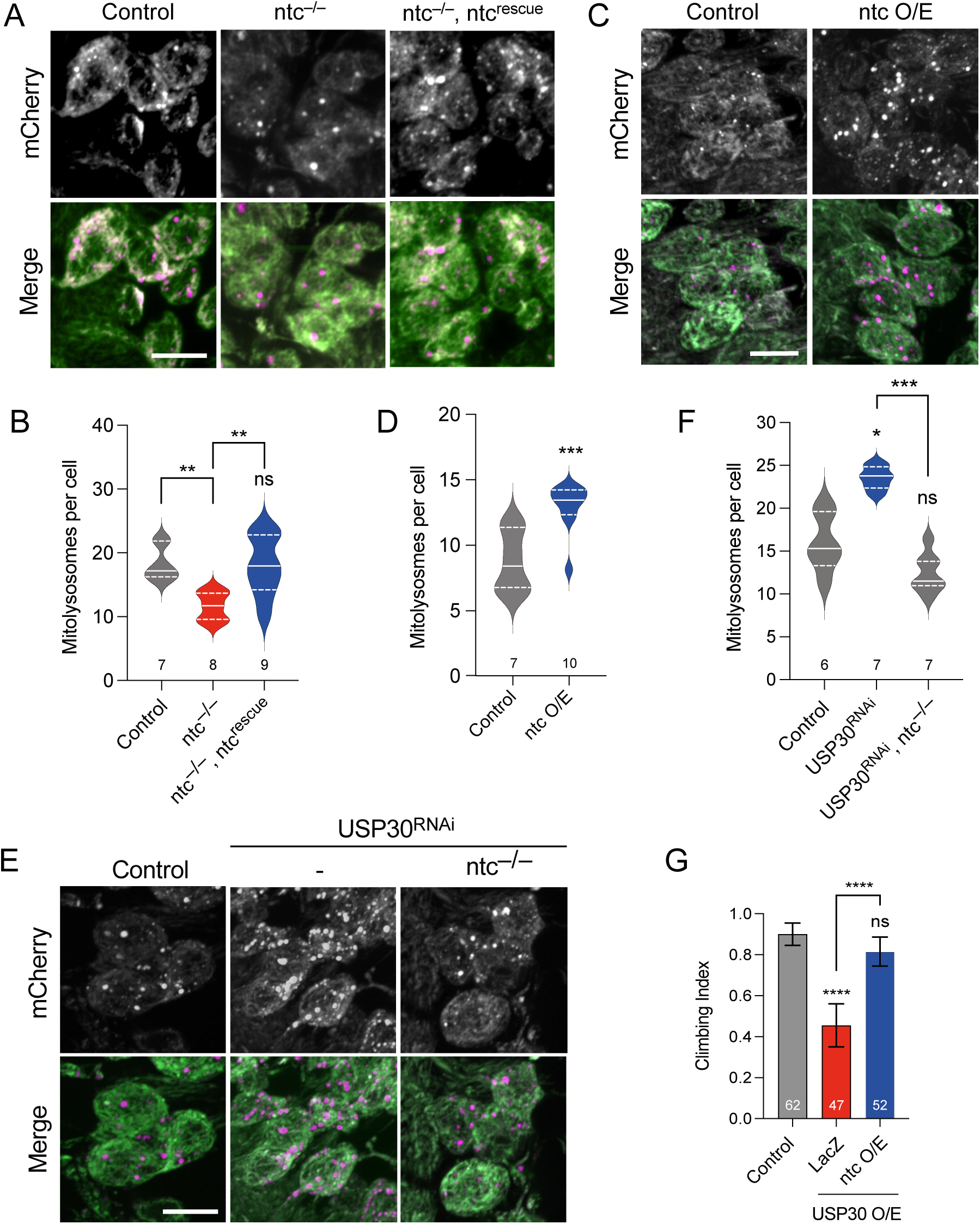
ntc affects basal mitophagy and counteracts USP30. (**A**, **B**) Confocal microscopy analysis of the *mito-QC* reporter in larval CNS of control and *ntc* mutant alone or with transgenic expression of *ntc* with the pan-neuronal driver *nSyb*-GAL4. Mitolysosomes are evident as GFP-negative/mCherry-positive (red-only) puncta. n shown in chart. One-way ANOVA with Bonferroni post-hoc test correction; *\*\* P* < 0.01. Scale bars = 10 μm. (**C**, **D**) Confocal microscopy analysis of the *mito-QC* reporter in larval CNS of control and transgenic expression of *ntc* with the pan-neuronal driver *nSyb*-GAL4. Mitolysosomes are evident as GFP-negative/mCherry-positive (red-only) puncta. n shown in chart. One-way ANOVA with Bonferroni post-hoc test correction; *\*\*\* P* < 0.001. Scale bars = 10 μm. (**E**, **F**) Confocal microscopy analysis of the *mito-QC* reporter in larval CNS of control and knockdown of USP30 with the pan-neuronal driver *nSyb*-GAL4 alone or in combination with *ntc* mutant. Mitolysosomes are evident as GFP-negative/mCherry-positive (red-only) puncta. n shown in chart. One-way ANOVA with Bonferroni post-hoc test correction; * *P* < 0.05, *\*\*\* P* < 0.001. Scale bars = 10 μm. (**G**) Climbing ability of 10-day-old flies overexpressing USP30 alone or in combination with ntc or parkin with the ubiquitous driver *Act*-GAL4. Chart show mean ± 95% CI; n shown in chart. Kruskal-Wallis nonparametric test with Dunn’s post-hoc test correction for multiple comparisons; **** *P* < 0.0001.

While stress-induced mitophagy has been extensively studied, especially in cultured cell lines, mechanisms regulating basal mitophagy are poorly characterised *in vivo*. Under basal conditions loss of *USP30*, an established regulator of mitophagy, leads to accumulation of ubiquitinated OMM proteins, and sustained *USP30* loss or inhibition promotes mitophagy *in vitro* (38–42), but *in vivo* characterisation remains limited. Upon *USP30* knockdown we observed a significant increase in basal mitophagy in larval neurons, using both the *mito-QC* (Figure 3E, 3F) and *mtx-QC* reporters (Figure S1A, S1B), and in adult flight muscle (Figure S3A, S3B). These results indicate that USP30 indeed inhibits basal mitophagy *in vivo* as expected. Importantly, we found that the induction of basal mitophagy by loss of *USP30* requires the activity of ntc as it was abolished in a *ntc* mutant background (Figure 3E, 3F).

As an orthogonal approach to test for genetic interaction at the organismal level, we found that while *USP30* knockdown did not cause any gross defect in adult viability or behaviour (Figure S3C), *USP30* overexpression caused a locomotor defect (Figure 3G). This phenotype was completely suppressed by co-expression of *ntc* (Figure 3G), consistent with the opposing actions of Ub-ligase and deubiquitinase and supporting the idea of ntc acting upstream of USP30. Interestingly, while *parkin* overexpression also suppressed the *USP30* overexpression phenotype as expected (Figure S3D), other Ub-ligases linked to mitophagy, Mul1 and March5 (39, 43–47), showed no suppression of this phenotype (Figure S3D). Moreover, similar to *parkin* loss (20) or overexpression (Figure S3E, S3F), loss of Mul1 or March5 had no effect on basal mitophagy (Figure S3G-S3J). Together, these data highlight the specificity of ntc and USP30 as important and opposing regulators of basal mitophagy.

Given the previously established links between FBXO7 and USP30 with toxin-induced PINK1/Parkin mitophagy in cultured human cells, we next analysed whether the induction of basal mitophagy by *ntc* overexpression or *USP30* knockdown involved the *Pink1/parkin* pathway *in vivo*. As previously reported (20), the absence of *parkin* or *Pink1* did not impact basal mitophagy in larval neurons (Fig 4A-D). However, overexpression of *ntc* was still able to induce mitophagy in the absence of either *parkin* or *Pink1* (Figure 4A-4D). Likewise, *USP30* knockdown also increased mitophagy independently of *parkin* and *Pink1* (Figure 5A-5D), consistent with *in vitro* findings (38, 39, 48, 49). Notably, the induction of mitophagy by *USP30* knockdown in *Pink1* or *parkin* mutant backgrounds required the function of *ntc* (Fig 5A-D), further underscoring the importance of ntc in basal mitophagy.

**Figure 4.**
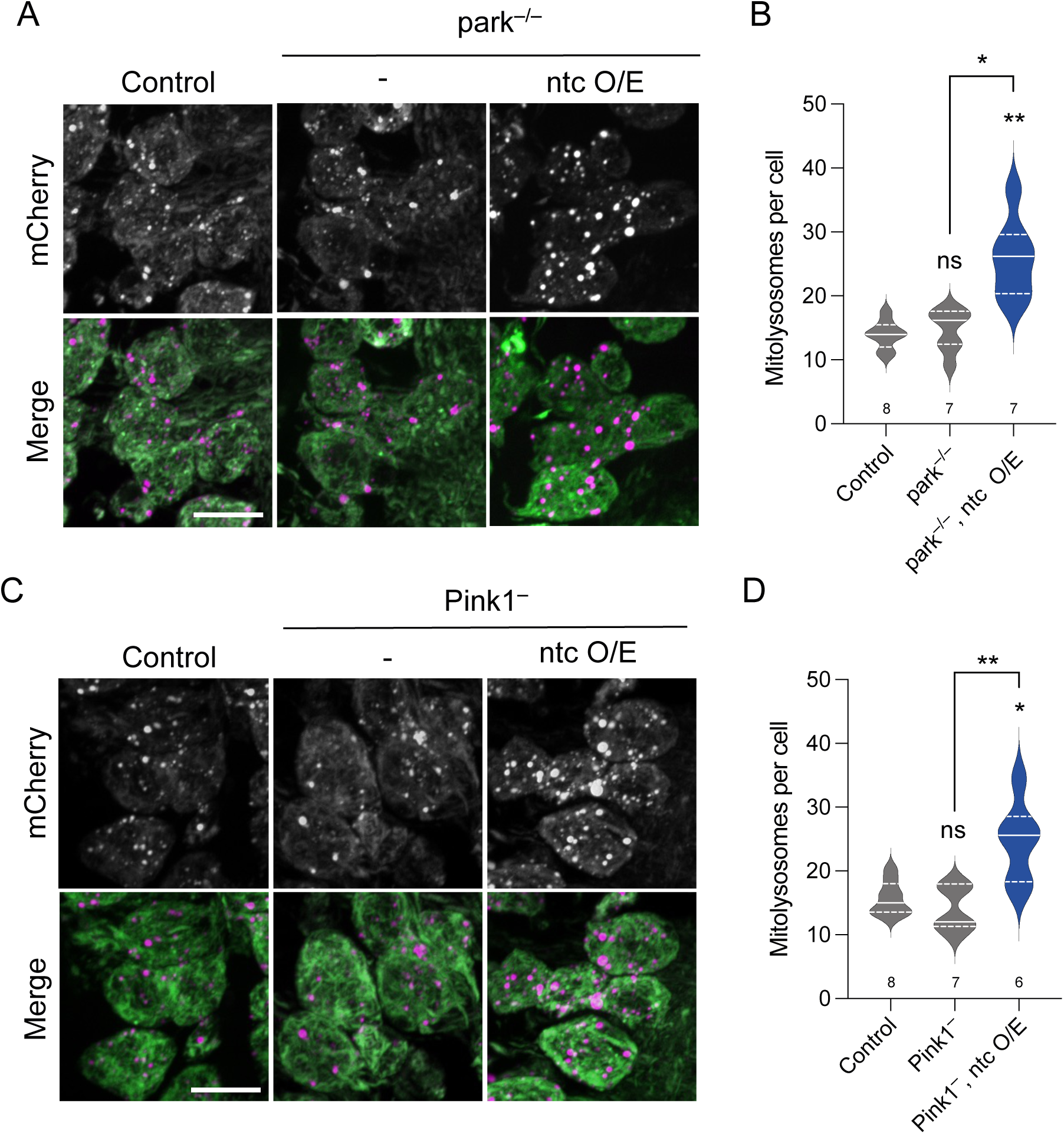
ntc regulates basal mitophagy in a parkin and Pink1 independent manner. (**A**, **B**) Confocal microscopy analysis of the *mito-QC* reporter in larval CNS of control and *parkin* mutant alone or with transgenic expression of *ntc* with the pan-neuronal driver *nSyb*-GAL4. Mitolysosomes are evident as GFP-negative/mCherry-positive (red-only) puncta. n shown in chart. One-way ANOVA with Bonferroni post-hoc test correction; * *P* < 0.05, *\*\* P* < 0.01. Scale bars = 10 μm. (**C**, **D**) Confocal microscopy analysis of the *mito-QC* reporter in larval CNS of control and *Pink1* mutant alone or with transgenic expression of *ntc* with the pan-neuronal driver *nSyb*-GAL4. Mitolysosomes are evident as GFP-negative/mCherry-positive (red-only) puncta. n shown in chart. One-way ANOVA with Bonferroni post-hoc test correction; * *P* < 0.05, *\*\* P* < 0.01. Scale bar = 10 μm.

**Figure 5.**
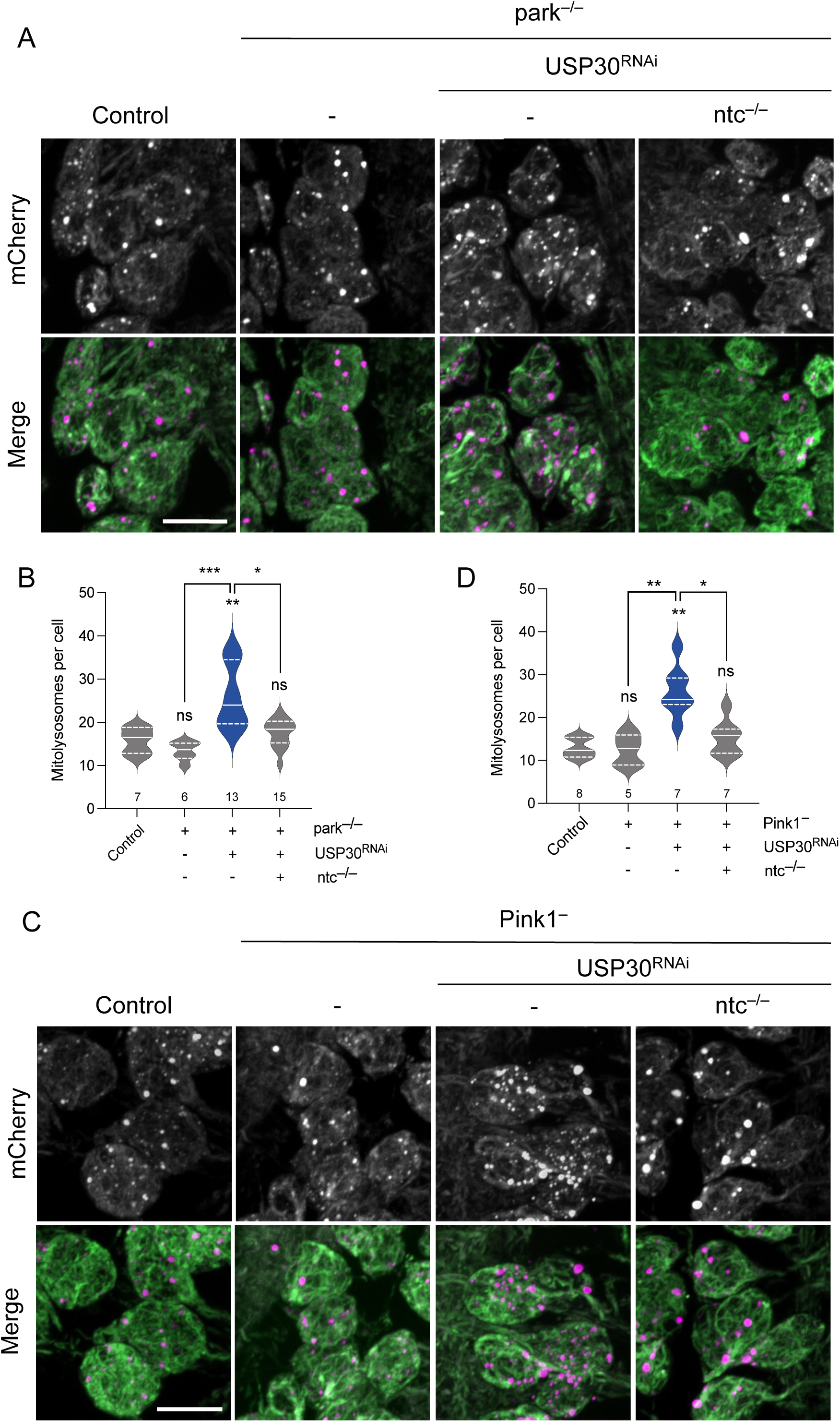
ntc is required for USP30 knockdown induced basal mitophagy in the absence of parkin or Pink1. (**A**, **B**) Confocal microscopy analysis of the mito-QC reporter in larval CNS of control, knockdown of USP30 with the pan-neuronal driver *nSyb*-GAL4 alone or in combination with *ntc* mutants in a *parkin* mutant background. Mitolysosomes are evident as GFP-negative/mCherry-positive (red-only) puncta. n shown in chart. One-way ANOVA with Bonferroni post-hoc test correction; * *P* < 0.05, *\*\* P* < 0.01, *\*\*\* P* < 0.001. Scale bars = 10 μm. (**C**, **D**) Confocal microscopy analysis of the *mito-QC* reporter in larval CNS of control, knockdown of USP30 with the pan-neuronal driver *nSyb*-GAL4 alone or in combination with *ntc* mutants in a *Pink1* mutant background. Mitolysosomes are evident as GFP-negative/mCherry-positive (red-only) puncta. n shown in chart. One-way ANOVA with Bonferroni post-hoc test correction; * *P* < 0.05, *\*\* P* < 0.01. Scale bar = 10 μm.

### General autophagy is not grossly affected upon USP30 or ntc manipulations

Although previously validated as a reliable mitophagy reporter *in vivo*, we nevertheless verified that the increased mitolysosome formation (mitophagy) by *ntc* overexpression or *USP30* knockdown occurs via canonical autophagy flux as it was abrogated in the absence of *Atg8a*, the main fly homologue of human ATG8 family members GABARAP/LC3, in both wild-type or *parkin* mutant backgrounds (Figure S4A-S4D). Notwithstanding, the observed increase in mitolysosomes could be due to an increase in non-selective autophagy (50, 51). Thus, we used a number of well-established orthogonal assays to assess the effect of these mitophagy-inducing manipulations on general autophagy.

First, we examined the level of lipidated Atg8a (Atg8a-II) which is incorporated into autophagosomal membranes and is used as an indication of autophagy induction (52, 53). Neither *USP30* knockdown, *ntc* overexpression nor *ntc* loss had any effect on autophagy induction (Figure 6A, 6B). Similarly, the steady-state levels of ref(2)P, the homologue of mammalian p62, which accumulates upon autophagic blockage (54), was also not affected in any of the conditions (Figure 6C, 6D). We also analysed the autophagy flux reporter GFP-mCherry-Atg8a (55–57). Quantification of mCherry-Atg8a puncta (autolysosomes) showed no changes upon manipulating levels of *USP30* or *ntc* (Figure 6E, 6F). Together, these results support that the effect observed upon manipulation of *USP30* or *ntc* is specific for basal mitophagy and not a consequence of altered general autophagic flux.

**Figure 6.**
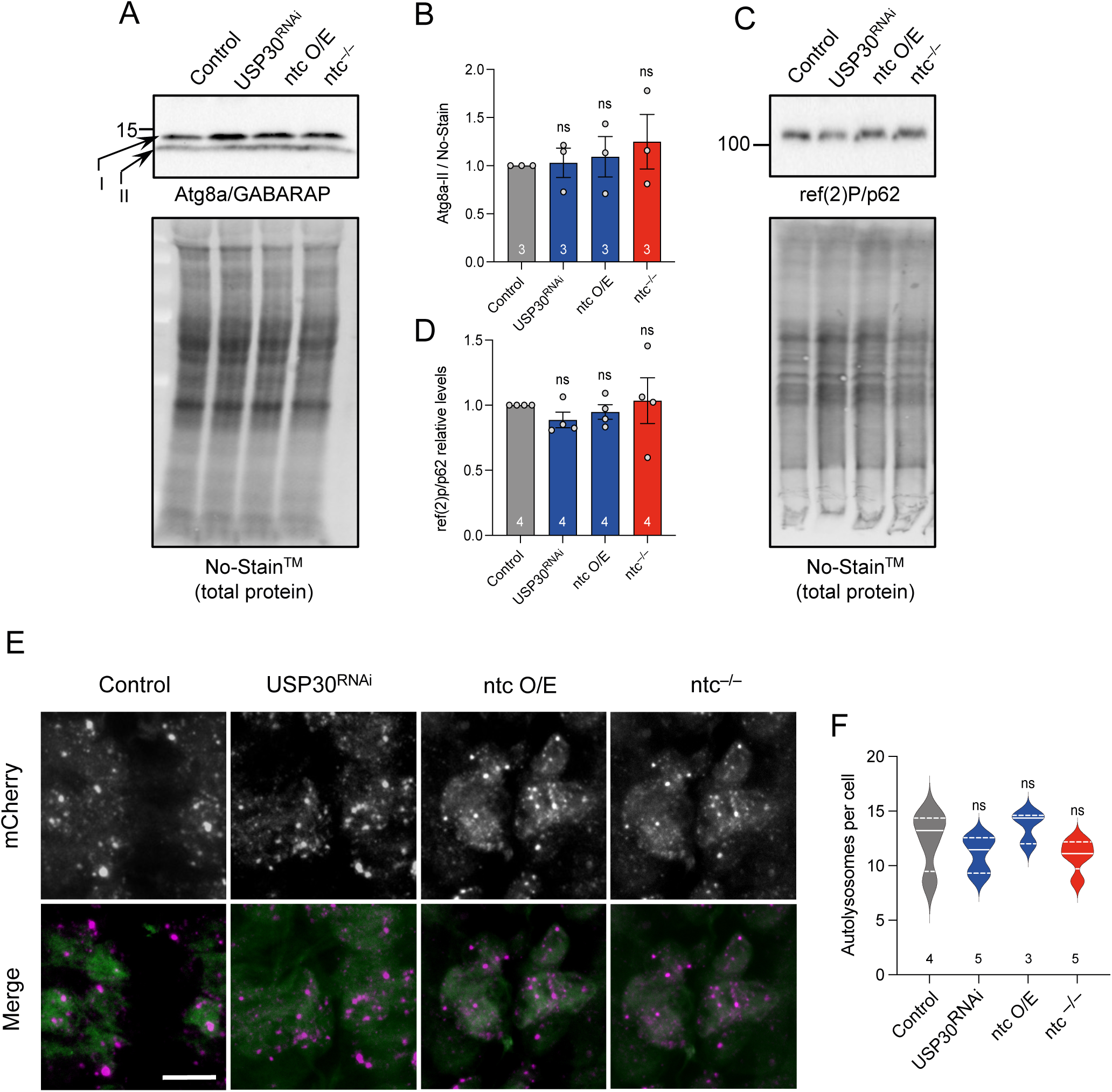
Manipulating mitophagy by altering levels of USP30 or ntc do not have an effect on general autophagy. (**A**, **B**) Representative immunoblot from whole fly lysates of Atg8a (non-lipidated) I and II (lipidated) in control, USP30 knockdown, ntc overexpression and *ntc* mutant with the ubiquitous driver *da*-GAL4. Data represented as mean ± SD; n shown in chart. One-way ANOVA with Bonferroni post-hoc test correction. (**C**, **D**) Representative immunoblot from whole fly lysates of ref(2)P/p62 in control, USP30 knockdown, ntc overexpression and *ntc* mutant with the ubiquitous driver *da*-GAL4. Data represented as mean ± SD; n shown in chart. One-way ANOVA with Bonferroni post-hoc test correction. (**E**, **F**) Confocal microscopy analysis and quantification of the red-only (autolysosomes) puncta per cell of larval CNS expressing the autophagy flux marker GFP-mCherry-Atg8a in combination with USP30 knockdown, ntc overexpression and *ntc* mutant with the pan-neuronal driver *nSyb*-GAL4. Data represented as mean ± SEM; n shown in chart. One-way ANOVA with Bonferroni post-hoc test correction; Scale bar = 10 μm.

### ntc increases basal mitochondrial ubiquitin and promotes phospho-ubiquitin formation

To gain a mechanistic insight into the regulation of basal mitophagy by ntc and USP30, we analysed their impact on mitochondrial ubiquitination. Consistent with their molecular functions, both *ntc* overexpression and *USP30* knockdown increased the total amount of ubiquitin present in mitochondria-enriched fractions under basal conditions (Figure 7A-B, Figure S5A), while total Ub remained unchanged (Figure S5B). Indeed, recent studies indicate that USP30 acts to remove pre-existing OMM ubiquitin under basal conditions, influencing the threshold needed for Parkin activation (38, 58).

**Figure 7.**
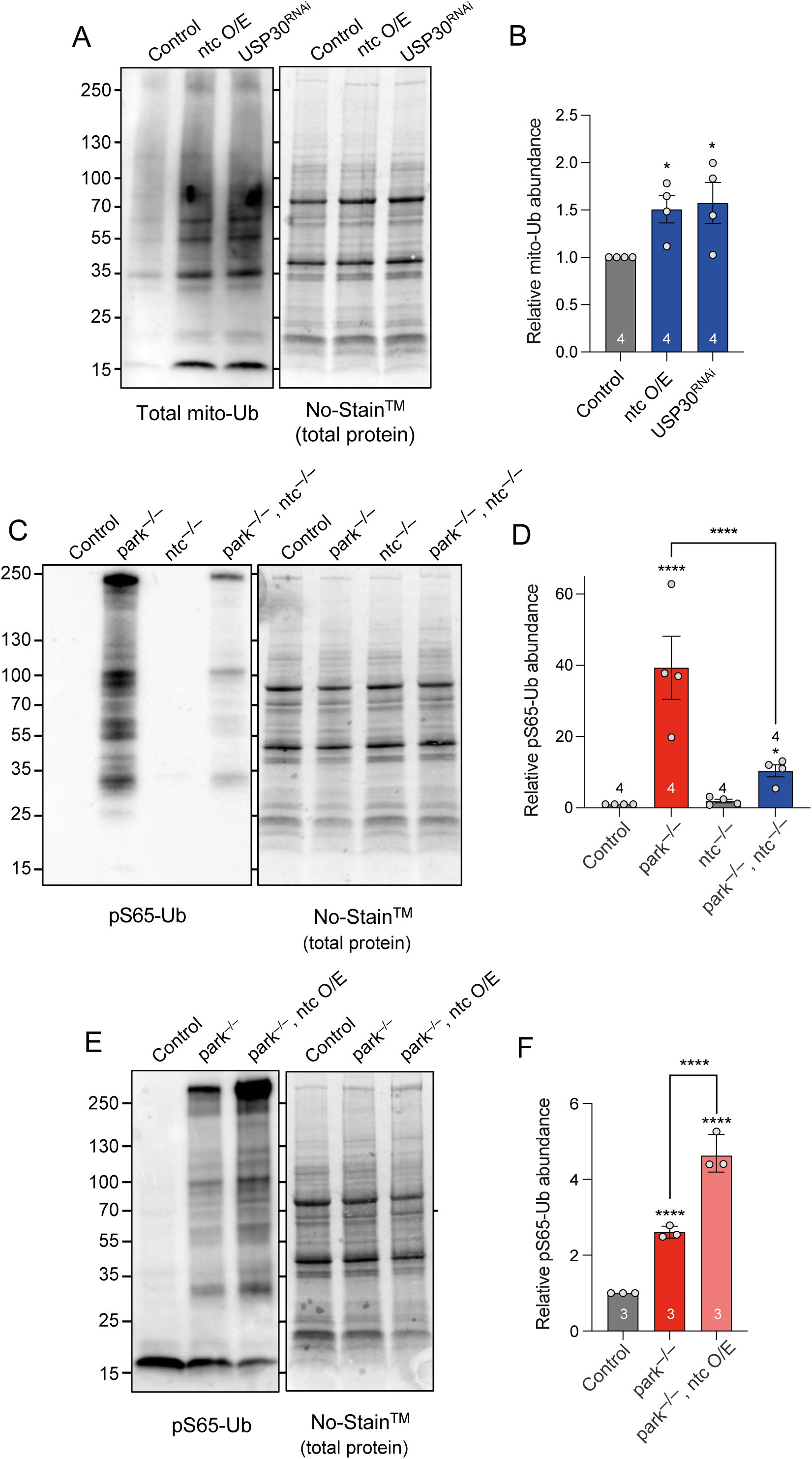
Both ntc overexpression and USP30 depletion promote accumulation of ubiquitin in the mitochondria and are necessary for pS65-Ub formation. (**A**) Representative immunoblot and quantification of (**B**) mitochondrial ubiquitin (P4D1) in control, ntc overexpression and USP30 knockdown with the ubiquitous driver *da*-GAL4. Data represented as mean ± SD; n shown in chart. One-way ANOVA with Bonferroni post-hoc test correction; * *P* < 0.05. (**C**, **D**) Representative immunoblot and quantification of mitochondrial pS65-Ub in *parkin* mutants, *ntc* mutants and the double *parkin*:*ntc* mutant. Data represented as mean ± SD; n shown in chart. One-way ANOVA with Bonferroni post-hoc test correction; * *P* < 0.05, **** *P* < 0.0001. (**E**, **F**) Representative immunoblot and quantification of mitochondrial pS65-Ub in control and *parkin* mutants alone or with the transgenic expression of ntc with the ubiquitous driver *da*-GAL4. Data represented as mean ± SD; n shown in chart. One-way ANOVA with Bonferroni post-hoc test correction; **** *P* < 0.0001.

We have recently described that loss of *parkin* causes a significant accumulation of pS65-Ub, consistent with parkin promoting the degradation of pS65-Ub labelled mitochondria (59). Unlike loss of *parkin*, loss of *ntc* alone did not lead to pS65-Ub build up (Figure 7C, 7D). However, *parkin*:*ntc* double mutants result in a significant reduction in the accumulated pS65-Ub levels (Figure 7C, 7D), and overexpression of *ntc* in the *parkin* mutant background increased the amount of pS65-Ub compared to *parkin* mutants alone (Figure 7E, 7F). These results are consistent with ntc promoting basal mitochondrial ubiquitination, which is phosphorylated by Pink1 to drive mitophagy. Thus, together these data suggest that ntc and USP30 work antagonistically to set up an OMM ubiquitin threshold needed for subsequent mitochondrial clearance.

## Discussion

Mutations in *FBXO7* have been linked to familial parkinsonism but the mechanisms of pathogenesis are poorly understood (22, 23). We have characterized the putative *Drosophila* homologue, *ntc*, to assess the functional homology with *FBXO7* and its potential as a model to understand the function of FBXO7 *in vivo*.

FBXO7 has been genetically linked to *Drosophila Pink1* and *parkin* and shown to facilitate PINK1/Parkin-mediated mitophagy *in vitro*. The *Drosophila* models have provided valuable insights into Pink1/parkin biology and genetic interactors have highlighted complementary pathways in mitochondrial homeostasis. Here we show that *ntc* mutants partially phenocopy *Pink1/parkin* phenotypes in locomotor function, lifespan and sensitivity to paraquat, consistent with a role in mitochondria quality control. We also observed the same genetic relationship as previously reported for *FBXO7* (27); namely, that overexpression of *ntc* can rescue all *parkin* mutant phenotypes but not for *Pink1* mutants. It is worth noting that in our previous study (27) we performed an initial characterisation of *ntc* as the putative homologue of *FBXO7* but observed no phenotypes reminiscent of *Pink1/parkin* mutants and provisionally concluded that ntc was not a functional homologue of FBXO7. However, these analyses were conducted using a hypomorphic allele (*ntc^f07259^*) in homozygosity, weakening the phenotype penetrance. In the current study, we have conducted our analyses with an amorphic allele, in parallel to studies comparing transgenic expression of *ntc* with *FBXO7*, hence supporting a functional homology between ntc and FBXO7.

Importantly, FBXO7 has previously been shown to facilitate the PINK1/Parkin pathway to promote stress-induced mitophagy *in vitro* (27, 60). However, whether it regulates mitophagy *in vivo* is poorly characterised. Indeed, mitophagy itself can occur via different regulators and stimuli, and represents only one of several mechanisms of mitochondrial quality control (61, 62). For instance, most studies have utilised a mitochondrial toxification stimulus to model substantial ‘damage’, however, basal mitophagy occurs as a housekeeping-type mechanism to regulate quantity or piecemeal turnover. The development of genetically encoded mitophagy reporters has provided a powerful approach to monitor both stress-induced or basal mitophagy *in vivo*. We have shown here that ntc is able to modulate basal mitophagy in a Pink1/parkin-independent manner, which is conserved by FBXO7 in human cell lines.

In comparison to stress-induced mitophagy, the regulation of basal mitophagy is poorly characterised, particularly in metazoa. While early studies showed the mitochondrial deubiquitinase USP30 antagonises PINK1/Parkin-mediated mitophagy (41, 63, 64), recent studies have established a role in basal mitophagy (38–40, 42). The emerging view suggests the main role of USP30 is acting as part of a quality control process for intra-mitochondrial proteins during import through the translocon (39, 42), and that basal mitochondrial ubiquitination provides the initiating substrate for PINK1 signalling. This paradigm requires a Ub ligase to provide basal ubiquitination that USP30 acts upon. While Phu et al. (42) suggested that this may be mediated via March5, this was not concurred by Ordureau et al. (39). Thus, the ligase providing the basal ubiquitination is unclear, and the role of USP30 in basal mitophagy *in vivo* remained to be established.

Here we demonstrated that knockdown of USP30 *in vivo* does indeed increase basal mitophagy in neurons and muscle, consistent with the *in vitro* studies. This also established a paradigm for a more deeply analysis of ntc’s function in basal mitophagy. While the induction of basal mitophagy by *USP30* knockdown is Pink1/parkin-independent as expected, it nevertheless required the function of ntc. In contrast, overexpression of parkin or depletion of either Mul1 or March5 did not affect neuronal basal mitophagy in our assay. The antagonistic relationship between USP30 and parkin was also evident from genetic interactions at the organismal level (rescue of a climbing defect), as expected, and was recapitulated with ntc. Such an interaction was also not evident with Mul1 and March5. Mechanistically, we found that *ntc* promotes mitochondrial ubiquitination under basal conditions, which was mirrored by the downregulation of *USP30*, consistent with the idea that basal ubiquitination signals basal mitophagy. Thus, our data suggest that ntc/FBXO7 provides basal ubiquitination, that is antagonised by USP30, which facilitates basal mitophagy upon a yet unclear stimulus.

The current mechanistic view of PINK1/Parkin stress-induced mitophagy signalling posits that upon mitochondrial depolarisation or import blocking, PINK1 becomes stabilised in the translocon, phosphorylates latent, pre-existing Ub, which promotes the recruitment of Parkin, triggering the feed-forward mechanism. Ubiquitination of import substrates would provide a ready source of latent Ub for PINK1 to phosphorylate as/when it becomes activated. Interestingly, USP30 has been found to be associated with TOMM20, a *bona fide* substrate of FBXO7 (65), and other translocon assemblies, where basally ubiquitinated translocon import substrates accumulate (39). Consistent with ntc providing the basal ubiquitination in this model, pS65-Ub that accumulates in *parkin* mutants is both reduced by loss of *ntc* and increased by its overexpression.

While we have shown that *ntc* overexpression is sufficient to induce basal mitophagy, independently of *Pink1/parkin*, and it is able to rescue *parkin* mutant phenotypes, curiously, it is not able to rescue *Pink1* phenotypes. As forced overexpression of *ntc* increases basal ubiquitination, it follows that already high levels of substrate for Pink1 circumvents the need for parkin ubiquitination activity (hence, why *ntc* overexpression can substitute for loss of *parkin*) when stress-induced mitophagy is required. However, despite high levels of latent ubiquitination, this alone is not sufficient to trigger stress-induced mitophagy in the absence of Pink1-mediated phosphorylation (hence, why *ntc* overexpression cannot rescue *Pink1* mutants). These findings further underscore the mechanistic and functional differences between basal and stress-induced mitophagy.

Analysing the potential role of ntc/FBXO7 in mitophagy and the relationship with Pink1/parkin *in vivo* has highlighted the complexity of how different forms of mitophagy may influence tissue homeostasis and relate to organismal phenotypes. As stated before, there are many different forms of mitochondrial quality control and mitophagy, and these perform different functions in cellular remodelling and homeostasis (61, 62). It follows that some forms of mitophagy may dramatically impact neuromuscular homeostasis when disrupted, while others may not. For instance, evidence supports that PINK1/Parkin promote stress-induced mitophagy but are minimally involved in basal mitophagy, and *Drosophila* mutants present multiple robust phenotypes. In contrast, ntc/FBXO7 (and USP30) regulate basal mitophagy and facilitate PINK1/Parkin mitophagy by providing the initiating ubiquitination, yet their mutant phenotypes only loosely resemble those of *Pink1/parkin* mutants.

How do these forms of mitophagy correlate with the mitochondrial/organismal phenotypes? The *Pink1/parkin* phenotypes are consistent with a catastrophic loss of integrity in energy-intensive, mitochondria-rich tissues caused by the lack of a crucial protective measure (induced mitophagy) for a specific circumstance (likely, mitochondrial ‘damage’ arising from a huge metabolic burst). In contrast, while loss of *ntc* causes a partial (but not complete) loss of basal mitophagy, the tissues affected in *Pink1/parkin* mutants appear to be able to cope with loss of this ‘housekeeping’ mitochondrial QC process, consistent with there being partially redundant pathways. Notably, these tissues are still able to mount a stress-induced response via Pink1/parkin as we have observed paraquat-induced accumulation of pS65-Ub in the *ntc* mutant (Figure S5C), reinforcing that stress-induced mitophagy can happen in the absence of ntc. Consequently, *ntc* mutants do not present the same degenerative phenotypes as *Pink1/parkin* mutants and maintain relatively normal mitochondrial function. Hence, it seems likely that the additional *ntc* mutant phenotypes of motor deficits and drastically short lifespan are due to additional functions of ntc/FBXO7, such as regulation of proteasome function and caspase activation, and not mitophagy.

In summary, we propose a model (Figure 8) in which FBXO7/ntc acts to prime OMM proteins with ubiquitin that is counteracted by USP30. This provides a key regulatory checkpoint in quality control, for instance, during mitochondrial protein import. It is likely that occasional defects in import or protein misfolding may lead to an increase of Ub above a threshold that is sufficient to provoke basal mitophagy. However, when the need arises this population of latent Ub could subsequently be phosphorylated by PINK1 (and amplified by Parkin) to trigger selective degradation upon specific stimulation. Although more studies are required to understand the interplay between FBXO7/ntc, USP30, Pink1 and parkin, this study provides a foundation to further elucidate the interplay between these mechanisms of mitochondrial quality control and reinforces its potential as a therapeutic target.

**Figure 8.**
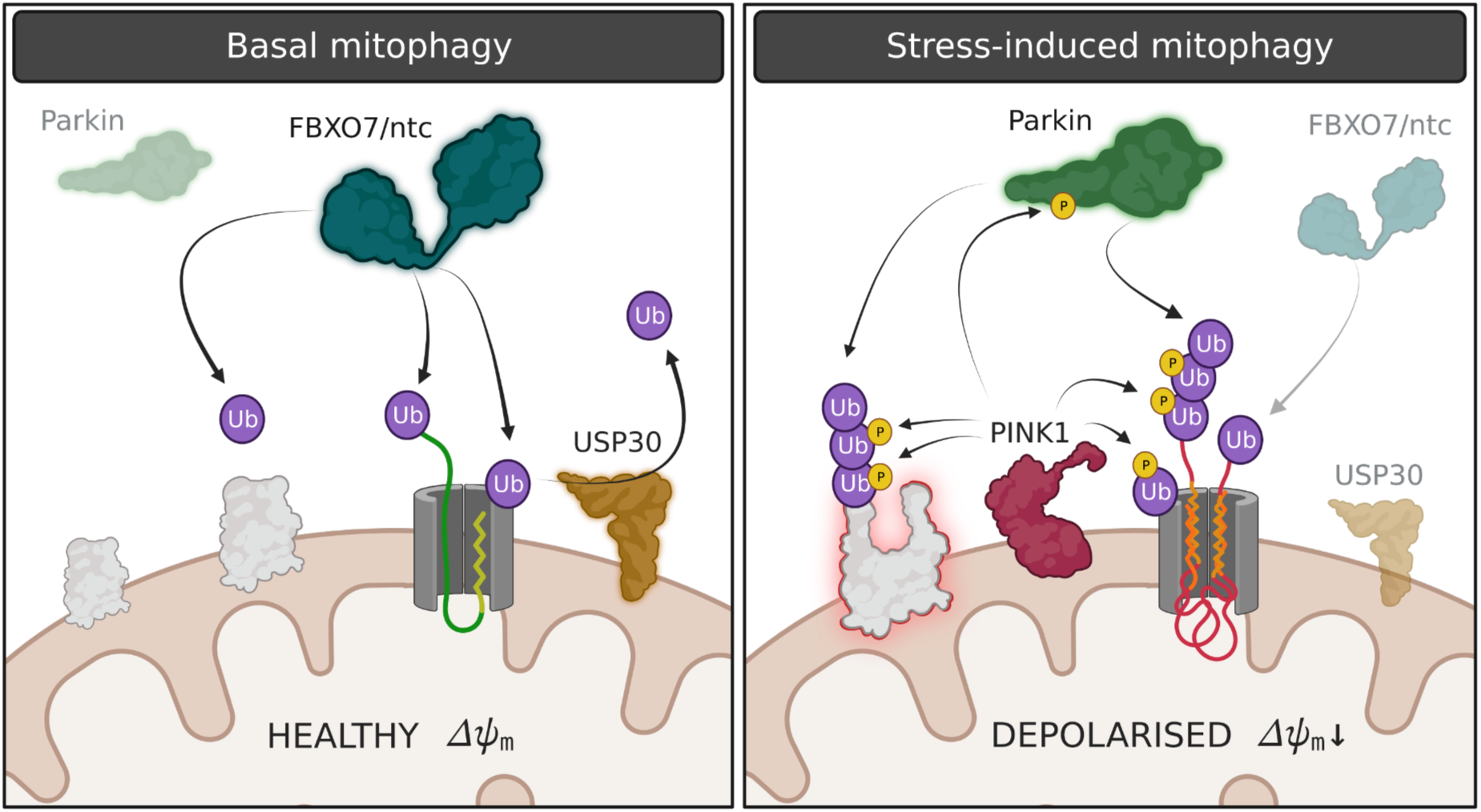
Proposed model for basal and stress-induced mitophagy regulated by ntc, USP30 and parkin. In basal, healthy conditions, FBXO7/ntc acts to prime OMM proteins with ubiquitin counteracted by USP30. This provides a key surveillance checkpoint in quality control of mitochondrial protein import and/or protein damage. Occasional defects in import or protein misfolding may lead to a Ub threshold that provokes basal mitophagy. However, when the need arises in the presence of a stress, for instance, where the membrane may become depolarised, this population of latent Ub could subsequently be phosphorylated by Pink1 and amplified by parkin to trigger selective degradation upon specific stimulation. Created with BioRender.com.

## Materials and Methods

### *Drosophila* stocks and procedures

*Drosophila* were raised under standard conditions in a temperature-controlled incubator with a 12h:12h light:dark cycle at 25 °C and 65 % relative humidity, on food consisting of agar, cornmeal, molasses, propionic acid and yeast. *ntc^ms771^*, *UAS-ntc* and *PBac{WH}CG10855^f07259^* strains were kindly provided by H. Steller. *park^25^* mutants and *UAS-parkin_C2_* (13), *UAS-FBXO7* (27), and *UAS-mito-QC* lines (20) have been previously described. *Pink1^B9^*mutants (12) were provided by J. Chung (SNU). *UAS-Mul1* and *Mul1^A6^*mutants were kindly provided by Ming Guo. *UAS-USP30* lines were kindly provided by Ugo Mayor. *UAS-USP30* RNAi (NIG-Fly 3016R[II]) was obtained from the NIG-Fly collection. The following strains were obtained from Bloomington *Drosophila* Stock Center (BDSC, RRID:SCR_006457): *w*^1118^ (RRID:BDSC_6326), *da-GAL4* (RRID:BDSC_55850), *nSyb-GAL4* (RRID:BDSC_51635), *GMR*-GAL4 (RRID:BDSC_1104), *Mef2*-GAL4 (RRID:BDSC_27390), *Act*-GAL4 (RRID:BDSC_25374), *Atg8a* [KG07569] *(Atg8a^−/−^)* (RRID:BDSC_14639), *UAS-GFP-mCherry-Atg8a* (RRID:BDSC_37749) and *Df(3L)Exel6097* (RRID:BDSC_7576). *UAS-lacZ* was obtained from FlyORF (RRID: FlyBase_FBst0503118). The following strains were obtained from Vienna Drosophila Research Centre (VDRC): *UAS-LacZ^RNAi^* (v51446), *UAS-March5* RNAi KK (v105711) and *UAS-March5* RNAi GD(v33309). *UAS-mtx-QC* and *UAS-March5* lines were generated as follows: *mtx-QC* mCherry-GFP was amplified from *UAS-mito-QC* and cloned in-frame with the mitochondrial targeting sequence from hCOX8A into pUAST.attB, and inserted in attP40 and attP16 sites; while for *March5* (*CG9855*), full length cDNA was cloned into pUAST vector and transgenesis performed by random insertion. A full description of the genotypes used in this study is shown in Supplementary table 1.

### Locomotor and lifespan assays

For locomotor assays, climbing (negative geotaxis assay) was assessed as previously described, with minor modifications (13). For lifespan experiments flies were grown under identical conditions at low-density. Progeny were collected under very light anaesthesia and kept in tubes of approximately 20 males each. Flies were transferred every 2-3 days to fresh tubes with normal food for normal lifespan and tubes with 10 mM paraquat in a filter paper for the oxidative stress lifespan. The number of dead flies was recorded on each transfer. Percent of survival was calculated at the end of the experiment after correcting for any accidental loss.

### Histology of adult thoraces

Thoraces were prepared from 5-day-old adult flies and treated as previously described (13). Semi-thin sections were then taken and stained with Toluidine blue, while ultra-thin sections were examined using a FEI Tecnai G2 Spirit 120KV transmission electron-microscope.

### Light microscopy imaging and scanning electron microscopy of *Drosophila* eye

Light microscopy imaging was assessed using a Nikon motorized SMZ stereo zoom microscope fitted with 1x Apo lens. Extended focus images were then generated using Nikon Elements software, using the same settings for all the genotypes. Flies were anaesthetised with CO2 during the process. Scanning electron microscopy (SEM) was performed according to a standard protocol (66). All animals of a given genotype displayed essentially identical phenotypes and randomly selected representative images are shown. Images were taken using a SEM microscope (Philips XL-20 SEM).

### Immunohistochemistry and sample preparation

*Drosophila* brains were dissected from 30-day-old flies and immunos-stained with anti-tyrosine hydroxylase (Immunostar Inc. #22491) as described previously (14). Brains were imaged with an Olympus FV1000 confocal with SIM-scanner on a BX61 upright microscope. Tyrosine hydroxylase-positive neurons were counted under blinded conditions. For immunostaining, adult flight muscles were dissected in PBS and fixed in 4% formaldehyde for 30 min, permeabilized in 0.3% Triton X-100 for 30 min, and blocked with 0.3% Triton X-100 plus 1% bovine serum albumin in PBS for 1 h at RT. Tissues were incubated with ATP5A antibody (Abcam Cat# ab14748, RRID:AB_301447; 1:500), diluted in 0.3% Triton X-100 plus 1% bovine serum albumin in PBS overnight at 4°C, then rinsed 3 times 10 min with 0.3% Triton X-100 in PBS, and incubated with the appropriate fluorescent secondary antibodies overnight at 4°C. The tissues were washed 3 times in PBS and mounted on slides using ProLong Diamond Antifade mounting medium (Thermo Fisher Scientific) and image next day. For mitolysosome analysis of *mito-QC* and *mtx-QC*, tissues were dissected and treated as previously described (20).

### Mitochondrial morphology in larval brain

Third instar larvae overexpressing *UAS-mitoGFP* with the pan-neuronal driver nSyb-GAL4 were dissected in PBS and fixed in 4% formaldehyde for 30 min. The tissues were washed 3 times in PBS and mounted on slides using ProLong Diamond Antifade mounting medium (Thermo Fisher Scientific) and image next day in a Zeiss LSM880 confocal microscope (63x / 1.4NA).

### ATP levels

The ATP assay was performed as described previously (67). Briefly, five male flies of the indicated age for each genotype were homogenized in 100 μL 6M guanidine-Tris/EDTA extraction buffer and subjected to rapid freezing in liquid nitrogen. Homogenates were diluted 1/100 with the extraction buffer and mixed with the luminescent solution (CellTiter-Glo Luminescent Cell Viability Assay (Promega, RRID:SCR_006724)). Luminescence was measured with a SpectraMax Gemini XPS luminometer (Molecular Devices). The average luminescent signal from technical triplicates was expressed relative to protein levels, quantified using the DC Protein Assay kit (Bio-Rad Laboratories, RRID:SCR_008426). Data from 2-4 independent experiments were averaged and the luminescence expressed as a percentage of the control.

### Respirometry

Mitochondrial respiration was assayed at 30 °C by high-resolution respirometry using an Oxygraph-2k high-resolution respirometer (OROBOROS Instruments) using a chamber volume set to 2 mL. Calibration with the air-saturated medium was performed daily. Data acquisition and analysis were carried out using Datlab software (OROBOROS Instruments). Five flies per genotype were homogenised in Respiration Buffer [120 mM sucrose, 50 mM KCl, 20 mM Tris–HCl, 4 mM KH_2_PO_4_, 2 mM MgCl_2_, and 1 mM EGTA, 1 g L^−1^ fatty acid-free BSA, pH 7.2]. For coupled (state 3) assays, complex I-linked respiration was measured at saturating concentrations of malate (2 mM), glutamate (10 mM), L-proline (10 mM) and adenosine diphosphate (ADP, 2.5 mM). Complex II-linked respiration was assayed in Respiration Buffer supplemented with 0.15 µM rotenone, 10 mM succinate and 2.5 mM ADP. Respiration was expressed as oxygen consumed per fly. Flies’ weight was equal in all genotypes tested.

### Image analysis and quantification of mitolysosomes

Analysis of mitolysosomes was done as previously described (20). Briefly, spinning disk microscopy–generated images from dissected larval brains or adult thoraces were processed using Imaris (version 9.0.2) analysis software (BitPlane; RRID:SCR_007370) to identify and count individual red-only puncta. The GFP and mCherry signals were adjusted to reduce background noise and retain only the distinct mitochondria network and red puncta, respectively. A surface rendered 3D structure corresponding to the mitochondria network was generated using the GFP signal. This volume was subtracted from the red channel to retain the mCherry signal that did not colocalize with the GFP-labelled mitochondria network. The mitolysosomes puncta were selected according to their intensity and an estimated size of 0.5 µm diameter, previously measured with Imaris. Additionally, the puncta were filtered with a minimum size cut off of 0.2 µm diameter. The remaining puncta were counted as the number of mitolysosomes. Larval CNS soma were analysed individually where discrete cells could be distinguished. The mean number of mitolysosomes per cell was calculated per animal. Data points in the quantification charts show mean mitolysosomes per cell for individual animals, where *n* ≥ 6 animals for each condition.

### Analysis and quantification of autolysosomes using the GFP-mCherry-Atg8a traffic light reporter

Third instar larval brains were dissected in phosphate buffered saline (PBS) and fixed with 4 % formaldehyde (FA) pH 7 (Thermo Scientific)/PBS for 20 minutes at room temperature. Samples were then washed in PBS followed by water to remove salts. ProLong antifade mounting media (Thermo Scientific) was used to mount the samples and imaged the day after. Confocal images, acquired with a Zeiss LSM880 microscope with the 63x/1.4 NA oil, were processed using FIJI (Image J). The quantification of autolysosomes was performed using FIJI (Image J) with the 3D Objects Counter Plugin. An area of interest was selected by choosing 6-10 cells per image. The threshold was based on matching the mask with the fluorescence. All puncta larger than 0.049 μm^3^ was considered an autolysosome. Data points in the quantification charts show mean mitolysosomes per cell for individual animals, where *n* ≥ 3 animals for each condition.

### Mitochondrial enrichment by differential centrifugation

All steps were performed on ice or at 4 °C. For immunoblotting analysis and biochemical fractionation from small numbers of flies (10–30), a modified mitochondrial enrichment procedure was performed. Flies were prepared either fresh or after flash-freezing in liquid nitrogen, with all direct comparisons performed with flies that were prepared in the same manner. Flies were transferred into a Dounce homogeniser containing 700 μL Solution A (70 mM sucrose, 20 mM HEPES pH 7.6, 220 mM mannitol, 1 mM EDTA) containing cOmplete protease inhibitors (Roche) and PhosSTOP phosphatase inhibitors (Roche), and manually homogenised with 50 strokes of a pestle. The homogenate was transferred to an Eppendorf tube, a further 500 μl of Solution A was added to the homogeniser and the sample was homogenised with a further 10 strokes. The homogenates were pooled and incubated for 30 minutes, then centrifuged for 5 minutes at 800 x g. The supernatant (containing mitochondria) was transferred to a new tube and clarified twice by centrifugation for 5 minutes at 1,000 x g. The clarified supernatant was then centrifuged for 10 minutes at 10,000 x g and the post-mitochondrial supernatant was discarded and the pellet retained for analysis. The mitochondrial pellet was washed once in Solution A containing only protease inhibitors, and then once in Solution A without inhibitors. The washed mitochondrial pellet was resuspended in 50 to 200 μL Sucrose Storage Buffer, the protein content determined by BCA assay (Thermo Pierce), and stored at −80 °C until needed.

### Antibodies and dyes

The following mouse antibodies were used for immunoblotting (WB) in this study: ATP5A (Abcam, ab14748, 1:10000), actin (Millipore, MAB1501, 1:5000), Ubiquitin (clone P4D1, Cell Signalling Technology, 3936, 1:1000), Ubiquitin (clone FK2, MBL, D058-3, 1:1000). The following rabbit antibodies were used in this study: pS65-Ub (Cell Signalling Technologies, 62802S, 1:1000), Marf ((16), 1:1000), GABARAP/Atg8a (Abcam ab109364, 1:1000), ref(2)P/p62 (Abcam ab178440, 1:1000). The following secondary antibodies were used: sheep anti-mouse (HRP-conjugated, GE Healthcare, NXA931V, 1:10000), donkey anti-rabbit (HRP-conjugated, GE Healthcare, NA934V, 1:10000).

### Whole-animal lysis and immunoblotting

For the analysis of whole cell lysates by immunoblot, 180 μL cold RIPA buffer (150 mM NaCl, 1% (v/v) NP-40, 0.5 % (w/v) sodium deoxycholate, 0.1% (w/v) SDS, 50 mM Tris pH 7.4), supplemented with cOmplete protease inhibitors, was added to 2 mL tubes containing 1.4 mm ceramic beads (Fisherbrand 15555799). Animals (5 to 20 per replicate) were harvested and stored on ice or flash-frozen in liquid N_2_, with all direct comparisons performed with flies that were harvested in the same manner. The flies were added to the tubes containing RIPA buffer and lysed using a Minilys homogeniser (Bertin Instruments) with the following settings: maximum speed, 10 seconds on, 10 seconds on ice, for a total of three cycles. After lysis, samples were returned to ice for 10 minutes then centrifuged 5 minutes at 21,000 x g, 4 °C. 90 μL supernatant was transferred to a fresh Eppendorf tube and centrifuged a further 10 minutes at 21,000 x g. 50 μL supernatant was then transferred to a fresh Eppendorf tube and the protein content determined by BCA assay as above. 30 μg total protein was then diluted in Laemmli Sample Buffer (Bio-Rad, 1610747) and analysed by SDS-PAGE using Mini-PROTEAN TGX Gels 4-20% (Bio-Rad, 4561093). For the analysis of mitochondria-enriched fractions, 30 μg mitochondrial protein was aliquoted into a tube, centrifuged 10 minutes at 16,000 x g, the supernatant removed and the pellet resuspended in Laemmli Sample Buffer (Bio-Rad, 1610747) to SDS-PAGE analysis as above. Gels were transferred onto pre-cut and soaked PVDF membranes (1704156, BioRad) using the BioRad Transblot Turbo transfer system, and blots were immediately stained with No-Stain™ Protein Labeling Reagent (Invitrogen, A44449) where indicated, according to the manufacturer’s instructions. Fluorescence intensity was measured using a BioRad Chemidoc MP using the IR680 setting. Blots were then washed by gentle shaking 3 times for 5 minutes in PBS containing 0.1% (v/v) Tween-20 (PBST), and blocked by incubation with PBST containing 3 % (w/v) BSA for 1 hour. Blots were washed a further 3 times as above then incubated at 4 °C overnight with primary antibodies in PBST containing 3 % (w/v) BSA. A further 3 washes were performed then the blots were incubated for one hour in secondary antibodies made up in PBST containing 3 % (w/v) BSA. Blots were then washed twice in PBST and once in PBS prior to incubation with ECL Prime western blotting System (Cytiva, RPN2232). Blots were imaged using the BioRad Chemidoc MP using exposure settings to minimise overexposure. Image analysis was performed using FIJI (Image J) and images were exported as TIFF files for publication. For paraquat treatment, flies were maintained in tubes (10-20 flies per replicate) containing 6 semi-circular pieces of filter paper (90 mm diameter, Cat#1001-090) saturated with 5% (w/v) sucrose solution containing 10 mM paraquat. Sucrose-only starvation experiments were performed as above, with the omission of paraquat. After 3 days, the flies were anaesthetised with mild CO_2_ and live flies only were harvested and process for immunoblotting as described above.

### ARPE-19-MQC cells

ARPE-19-MQC-FIS1 cells were used to assess mitophagy in a human cell model. Briefly, cells were transfected in reverse with RNAiMax (13778075, Thermo Fisher Scientific) and non-targeting control siRNA (ON-TARGETplus Non-targeting Control Pool, D-001810-10-20, Horizon Discovery) or FBXO7 siRNA (ON-TARGETplus Human FBXO7 siRNA, L-013606-00-0010, Horizon Discovery). After 48h knockdown cells were seeded into and ibidi dish (IB-81156) Thistle Scientific Ltd), and after 72h cells were imaged live by a spinning disk microscope and generated images were processed using Imaris as previously described.

## Statistical analysis

Data are reported as mean ± SEM or mean ± 95% CI as indicated in figure legends, and the n numbers of distinct biological replicates are shown in each graph. For statistical analyses of lifespan experiments, a log rank (Mantel-Cox) test was used. For behavioural analyses, a Kruskal-Wallis nonparametric test with Dunn’s post-hoc test correction for multiple comparisons was used. Where multiple groups were compared, statistical significance was calculated by one-way ANOVA with Bonferroni post-hoc test correction for multiple samples. This test was applied to dopaminergic neuron analysis and mitophagy where more than two groups were compared. For Ub and pS65-Ub abundance a one-way ANOVA with Dunnett’s correction for multiple comparisons was used. When only two groups were compared a Welch’s *t* test was used. The absence of statistical analysis in between any group reflects “no statistical significance”. All the samples were collected and processed simultaneously and therefore no randomization was appropriate. Unless otherwise indicated, all the acquisition of images and analysis was done in blind conditions. Statistical analyses were performed using GraphPad Prism 9 software (GraphPad Prism, RRID:SCR_002798). Statistical significance was represented in all cases as follows: *\* P* < 0.05, *\*\* P* < 0.01, *\*\*\* P* < 0.001 and **** *P* < 0.0001.

## Data Availability

All data needed to evaluate the conclusions in the paper are present in the paper and/or the Supplementary Materials. This study includes no data deposited in external repositories. Additional data related to this paper may be requested from the authors.

## Acknowledgements

We kindly thank Herman Steller for generously sharing the ntc lines, Ugo Mayor for the USP30 overexpression lines and Ian Ganley for the ARPE-19-MQC cells. We thank Roberta Tufi for performing the ATP assay, Wing Hei Au and Federica De Lazzari for comments and edits on the manuscript, and all the members of the Whitworth’s lab for discussions.

## Author contributions

A.S.M. conceived the project, designed the experiments, performed experimental work, analysed data, wrote the original draft, revised and edited the manuscript.

A.M. conceived parts of the project, designed and performed some experimental work, analysed data and revised and edited the manuscript.

A.J.W. conceived and supervised the project, wrote the original draft, revised and edited the manuscript.

## Figure Legends

**Figure S1.**
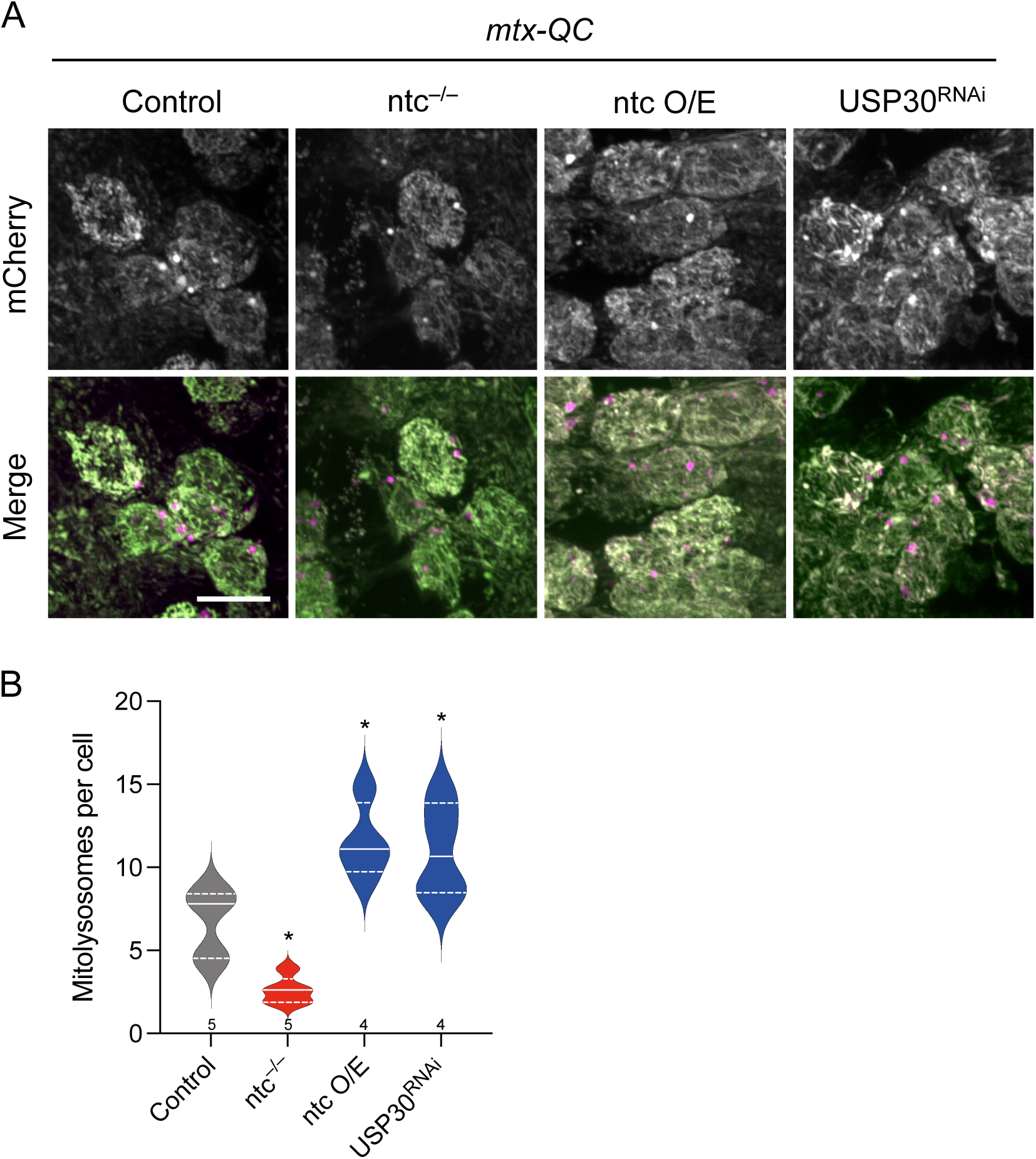
*mito-QC* reporter targeted to the mitochondrial matrix (*mtx-QC*) reproduces the results observed with the *mito-QC*. (**A**, **B**) Confocal microscopy analysis of the *mtx-QC* reporter in larval CNS of control, *ntc* mutant, *ntc* overexpression and USP30 knockdown with the pan-neuronal driver *nSyb*-GAL4. Mitolysosomes are evident as GFP-negative/mCherry-positive (red-only) puncta. n shown in chart. One-way ANOVA with Bonferroni post-hoc test correction; * *P* < 0.05. Scale bars = 10 μm.

**Figure S2.**
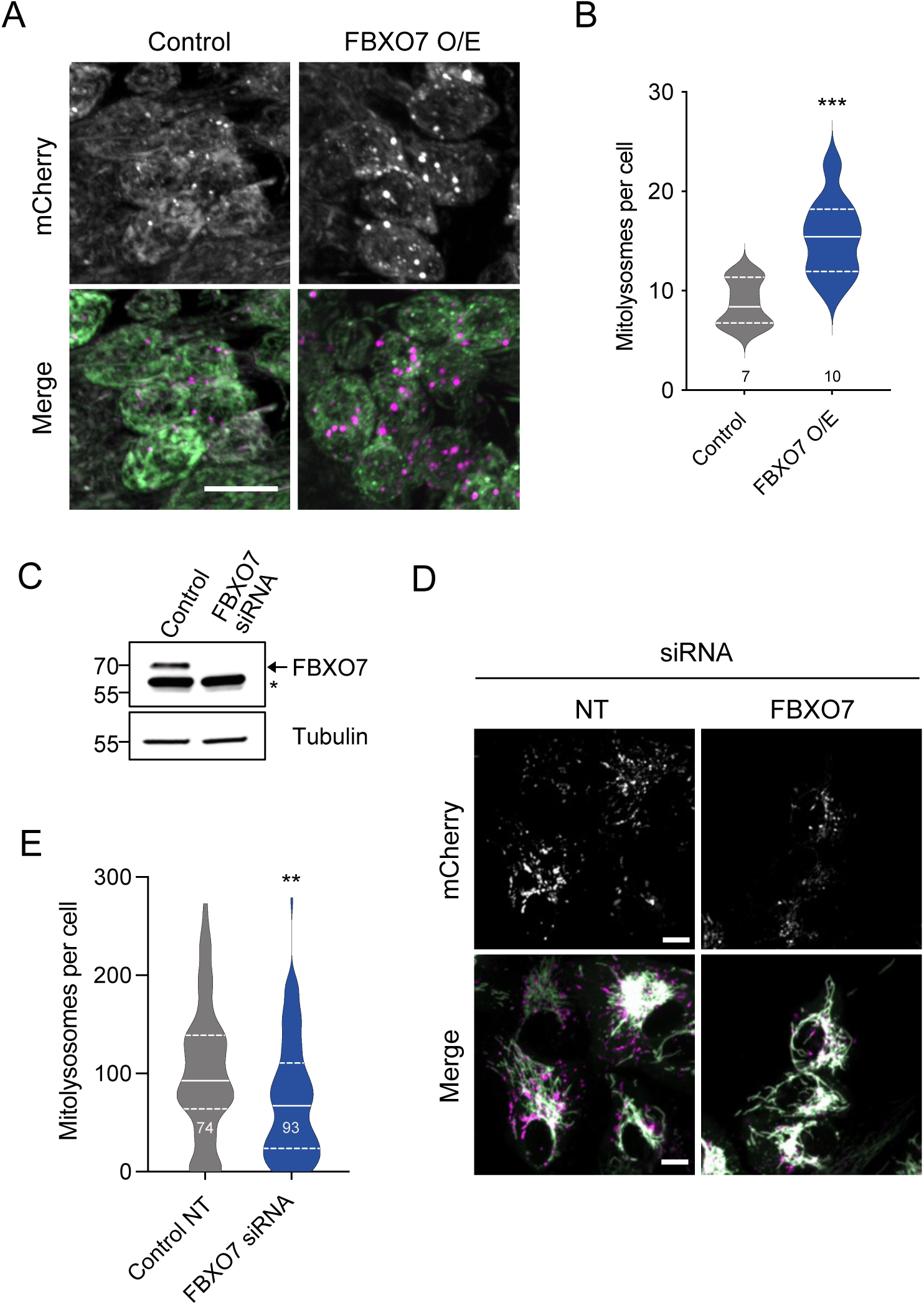
FBXO7 affects basal mitophagy *in vivo Drosophila* neurons and in a human cell line. (**A**, **B**) Confocal microscopy analysis of the *mito-QC* reporter in larval CNS of control and the transgenic expression of *FBXO7* with the pan-neuronal driver *nSyb*-GAL4. Mitolysosomes are evident as GFP-negative/mCherry-positive (red-only) puncta; n shown in chart. One-way ANOVA with Bonferroni post-hoc test correction; *\*\*\* P* < 0.001. Scale bar = 10 μm. (**C**) Immunoblot analysis of the knockdown of FBXO7 in human ARPE-19 cells expressing the *mito-QC* reporter. Arrow shows FBXO7 band; * shows non-specific band. (**D**, **E**) Confocal microscopy analysis of the *mito-QC* reporter in ARPE-19 human cell line of control siRNA and FBXO7 siRNA. Mitolysosomes are evident as GFP-negative/mCherry-positive (red-only) puncta. n shown in chart. Two-tailed *t* test; *\*\* P* < 0.01. Scale bars = 10 μm.

**Figure S3.**
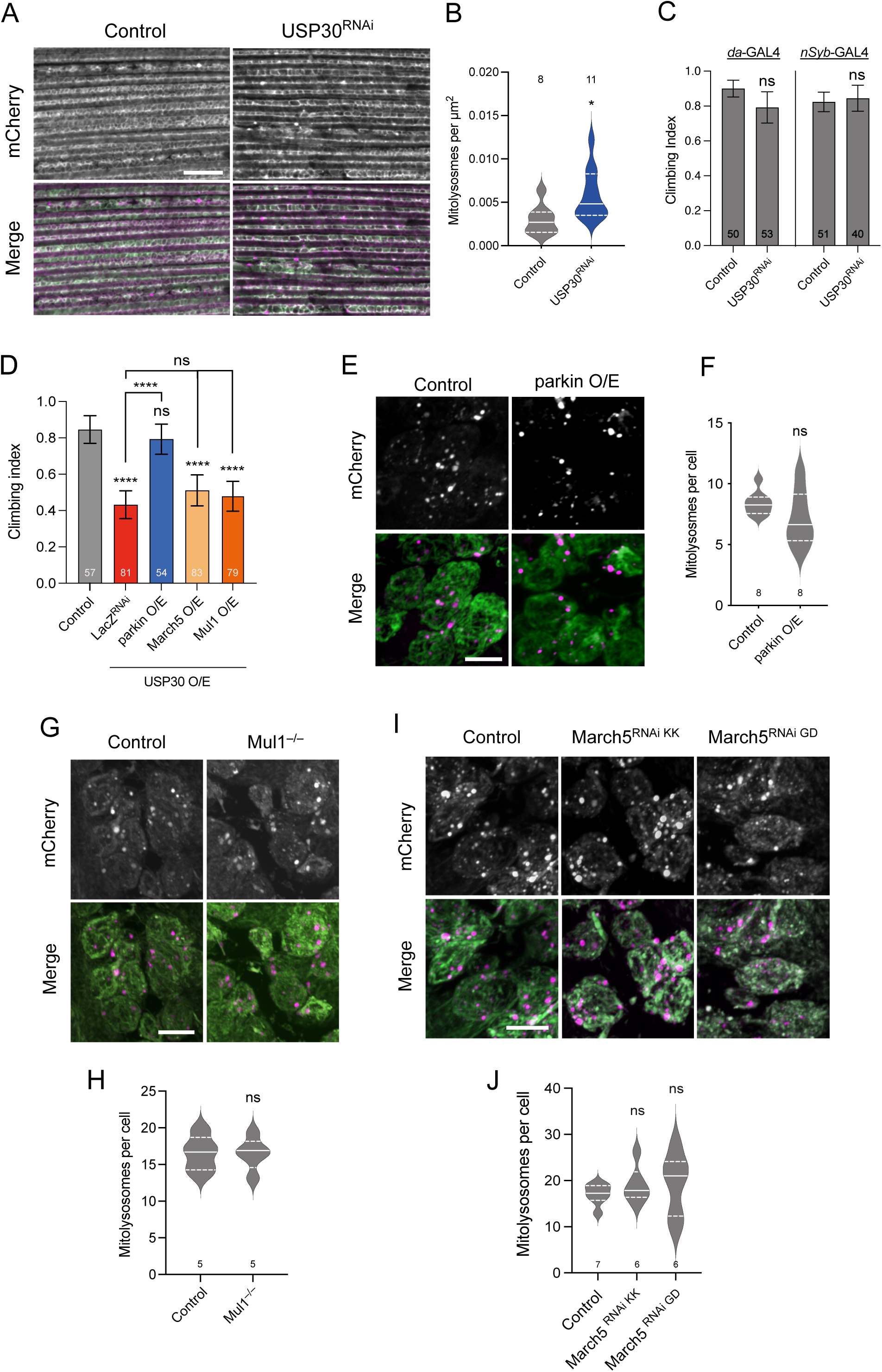
USP30 but neither March5 nor MUL1 can modulate mitophagy *in vivo*. (**A**, **B**) Confocal microscopy analysis of the *mito-QC* reporter in 2-day-old adult thoraces of control and USP30 knockdown with the muscular driver *Mef2*-GAL4. Mitolysosomes are evident as GFP-negative/mCherry-positive (red-only) puncta. n shown in chart. Two-tailed *t* test; * *P* < 0.05. Scale bar = 10 μm. (**C**) Climbing ability of 2-day-old flies expressing *USP30* RNAi with the ubiquitous driver *da*-GAL4 or with the pan-neuronal driver *nSyb*-GAL4. Chart show mean ± 95% CI and n values. Kruskal-Wallis nonparametric test with Dunn’s post-hoc test correction for multiple comparisons. (**D**) Climbing ability of 10-day-old flies overexpressing USP30 alone or in combination with parkin, March5 or Mul1 with the ubiquitous driver *Act*-GAL4. Chart show mean ± 95% CI and n values. Kruskal-Wallis nonparametric test with Dunn’s post-hoc test correction for multiple comparisons; **** *P* < 0.0001. (**E**, **F**) Confocal microscopy analysis of the *mito-QC* reporter in larval CNS of control and parkin overexpression with the pan-neuronal driver *nSyb*-GAL4. Mitolysosomes are evident as GFP-negative/mCherry-positive (red-only) puncta; n shown in chart. Two-tailed *t* test. Scale bar = 10 μm. (**G**, **H**) Confocal microscopy analysis of the *mito-QC* reporter in larval CNS of control and *Mul1* mutant. Mitolysosomes are evident as GFP-negative/mCherry-positive (red-only) puncta. n shown in chart. Two-tailed *t* test. Scale bar = 10 μm (**I**, **J**) Confocal microscopy analysis of the *mito-QC* reporter in larval CNS of control and March5 knockdowns with the pan-neuronal driver *nSyb*-GAL4. Mitolysosomes are evident as GFP-negative/mCherry-positive (red-only) puncta. n shown in chart. Two-tailed *t* test. Scale bar = 10 μm.

**Figure S4.**
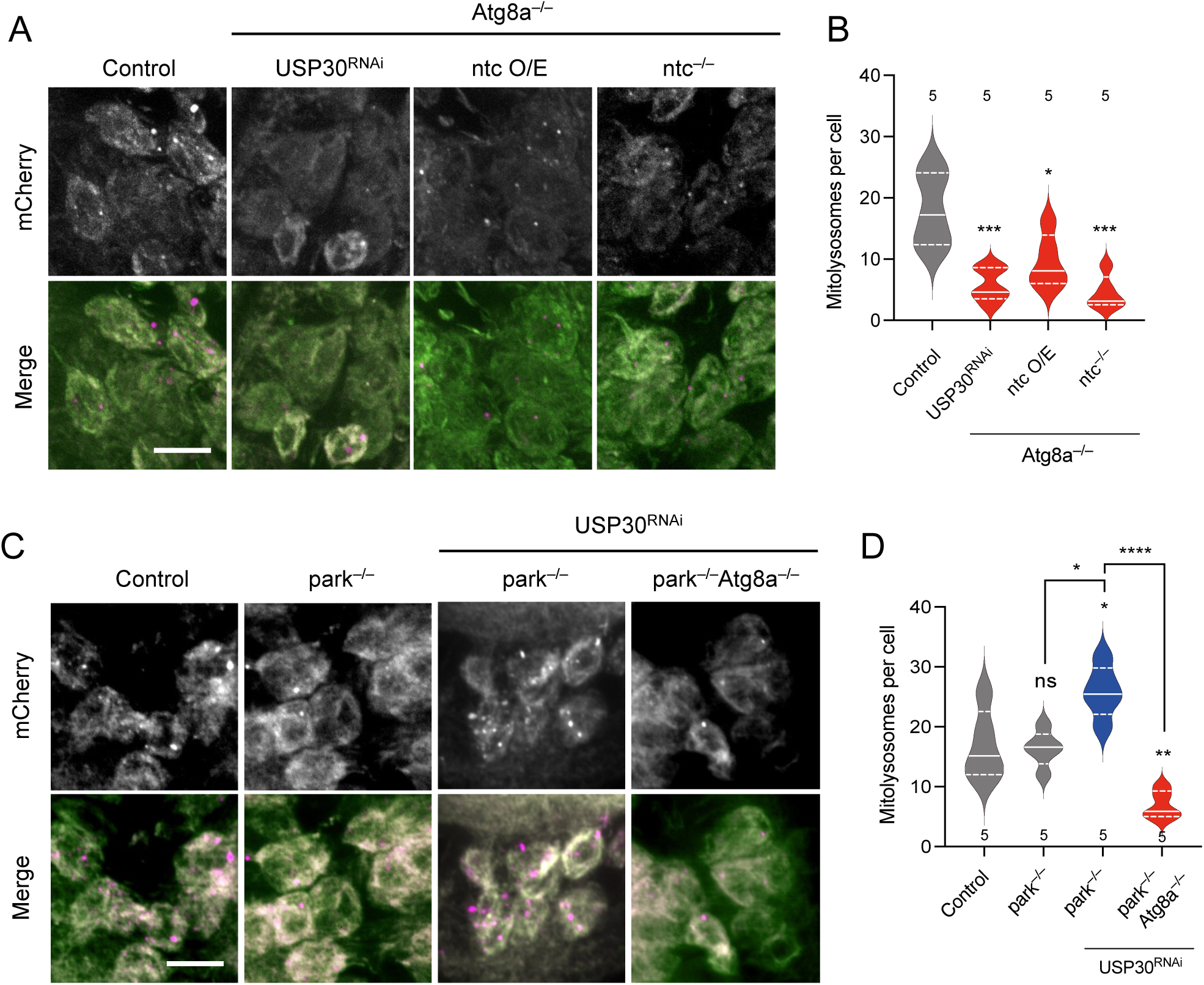
Increase in mitophagy levels requires the critical autophagy factor Atg8. (**A**, **B**) Confocal microscopy analysis of the *mito-QC* reporter in larval CNS of control, knockdown of USP30, overexpression of ntc and *ntc* mutant in the *Atg8a* mutant background with the pan-neuronal driver *nSyb*-GAL4. Mitolysosomes are evident as GFP-negative/mCherry-positive (red-only) puncta. n shown in chart. One-way ANOVA with Bonferroni post-hoc test correction; * *P* < 0.05, *\*\*\* P* < 0.001. Scale bar = 10 μm. (**C**, **D**) Confocal microscopy analysis of the *mito-QC* reporter in larval CNS of control and *parkin* mutant alone or with the USP30 knockdown in the presence or absence of *Atg8a*, with the pan-neuronal driver *nSyb*-GAL4. Mitolysosomes are evident as GFP-negative/mCherry-positive (red-only) puncta; n shown in chart. One-way ANOVA with Bonferroni post-hoc test correction; * *P* < 0.05, *\*\* P* < 0.01, **** *P* < 0.0001. Scale bar = 10 μm.

**Figure S5.**
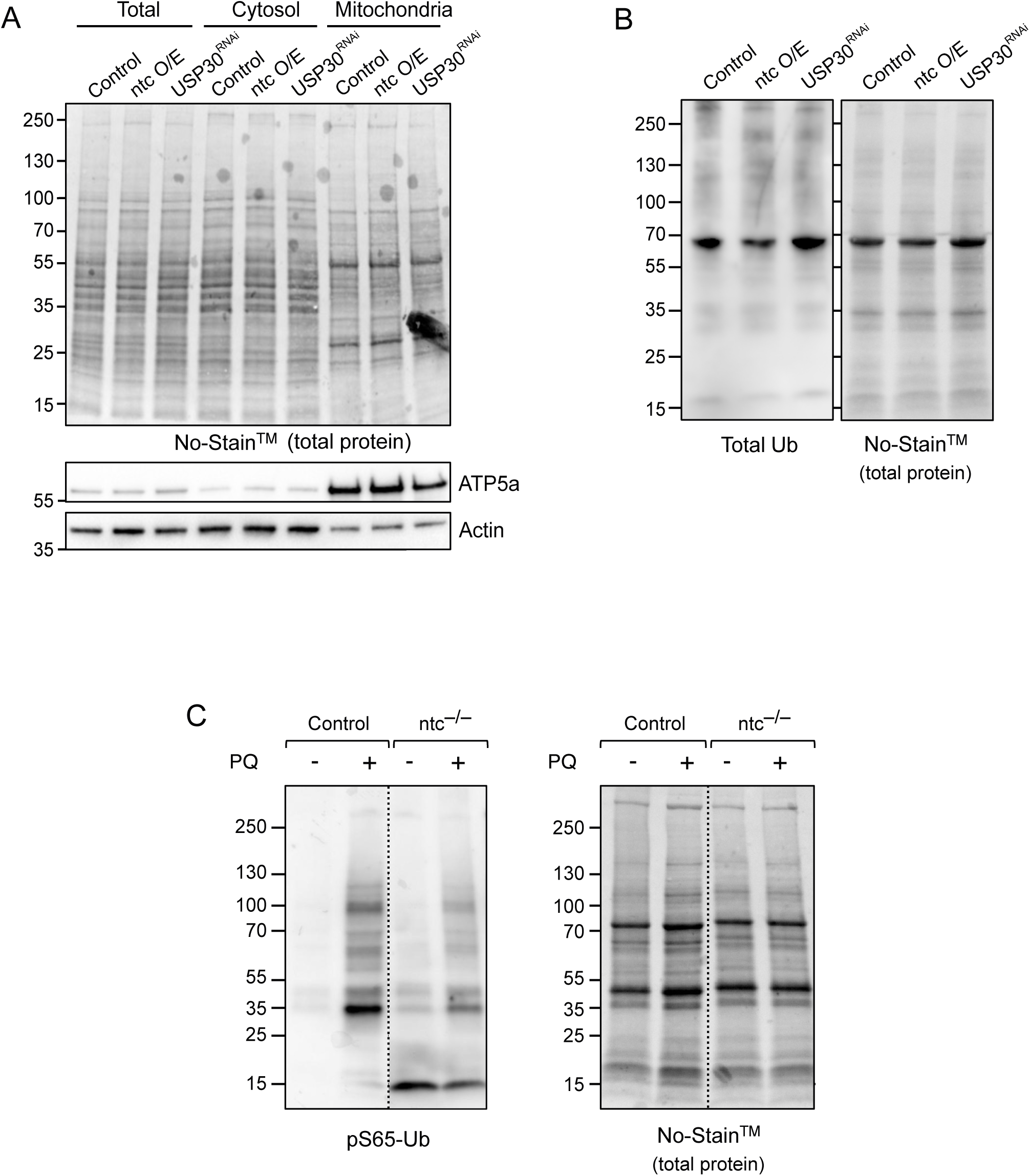
Analysis of total Ub in subcellular fractions and pS65-Ub upon oxidative stress-induction. (**A**) Representative immunoblot of the subcellular fractionation of 2-day-old flies with the transgenic expression of *ntc* or *USP30* RNAi with the ubiquitous driver *da*-GAL4. Cytosolic- and mitochondria-enriched fractions are label with Actin and ATP5a, respectively. **(B)** Representative immunoblot of total ubiquitin (FK2) of 2-day-old flies in control, *ntc* overexpression and *USP30* knockdown with the ubiquitous driver *da*-GAL4. **(C)** Representative immunoblot of pS65-Ub levels of 2-day-old flies treated with paraquat (PQ) in control and *ntc* mutants.

**Supplementary Table 1.** Details of full genotypes used in this study. More details of each line can be found in Methods.

| **Figure 1** |  |
| --- | --- |
| **<u>Label</u>** | **<u>Genotype</u>** |
| **A** |  |
| Control | UAS-LacZ/+; da-GAL4/+ |
| park<sup>-/-</sup> | UAS-LacZ/+; da-GAL4, park<sup>25</sup>/park<sup>25</sup> |
| park<sup>-/-</sup>; ntc O/E | UAS-ntc/+; da-GAL4, park<sup>25</sup>/park<sup>25</sup> |
| **B** |  |
| Control | UAS-LacZ/+; da-GAL4/+ |
| park<sup>-/-</sup> | UAS-LacZ/+; da-GAL4, park<sup>25</sup>/park<sup>25</sup> |
| park<sup>-/-</sup>; ntc O/E | UAS-ntc/+; da-GAL4, park<sup>25</sup>/park<sup>25</sup> |
| **C** |  |
| Control | UAS-LacZ/+; da-GAL4/+ |
| park<sup>-/-</sup> | UAS-LacZ/+; da-GAL4, park<sup>25</sup>/park<sup>25</sup> |
| park<sup>-/-</sup>; ntc | UAS-ntc/+; da-GAL4, park<sup>25</sup>/park<sup>25</sup> |
| **D** |  |
| Control | UAS-LacZ/+; da-GAL4/+ |
| park<sup>-/-</sup> | UAS-LacZ/+; da-GAL4, park<sup>25</sup>/park<sup>25</sup> |
| park<sup>-/-</sup>; ntc O/E | UAS-ntc/+; da-GAL4, park<sup>25</sup>/park<sup>25</sup> |
| **E** |  |
| Control | UAS-LacZ/+; da-GAL4/+ |
| park<sup>-/-</sup> | UAS-LacZ/+; da-GAL4, park<sup>25</sup>/park<sup>25</sup> |
| park<sup>-/-</sup>; ntc O/E | UAS-ntc/+; da-GAL4, park<sup>25</sup>/park<sup>25</sup> |
| **F** |  |
| Control | UAS-LacZ/+; da-GAL4/+ |
| Pink1<sup>-</sup> | Pink1<sup>B9</sup>/Y; UAS-LacZ/+; da-GAL4/+ |
| Pink1<sup>-</sup>; ntc O/E | Pink1<sup>B9</sup>/Y; UAS-ntc/+; da-GAL4/+ |
| **G** |  |
| Control | UAS-LacZ/+; da-GAL4/+ |
| Pink1<sup>-</sup> | Pink1<sup>B9</sup>/Y; UAS-LacZ/+; da-GAL4/+ |
| Pink1<sup>-</sup>; ntc O/E | Pink1<sup>B9</sup>/Y; UAS-ntc/+; da-GAL4/+ |
| **H** |  |
| Control | UAS-LacZ/+; da-GAL4/+ |
| Pink1<sup>-</sup> | Pink1<sup>B9</sup>/Y; UAS-LacZ/+; da-GAL4/+ |
| Pink1<sup>-</sup>; ntc O/E | Pink1<sup>B9</sup>/Y; UAS-ntc/+; da-GAL4/+ |
| **I** |  |
| Control | GMR-GAL4/UAS-LacZ |
| ntc O/E | GMR-GAL4, UAS-LacZ/UAS-ntc |
| Pink1 O/E | GMR-GAL4, UAS-Pink1/UAS-LacZ |
| Pink1+ntc O/E | GMR-GAL4, UAS-Pink1/UAS-ntc |

| **Figure 2** |  |
| --- | --- |
| <u>Label</u> | <u>Genotype</u> |
| **A** |  |
| Control | UAS-LacZ/+; da-GAL4/+ |
| ntc<sup>-/-</sup> | ntc<sup>ms771</sup>/ntc<sup>ms771</sup> |
| ntc<sup>-/Df</sup> | ntc<sup>ms771</sup>/Df(3L)Exel6097 |
| ntc<sup>-/-</sup>, ntc<sup>rescue</sup> | UAS-ntc; da-GAL4, ntc<sup>ms771</sup>/ ntc<sup>ms771</sup> |
| **B** |  |
| Control | UAS-LacZ/+; da-GAL4/+ |
| ntc<sup>-/-</sup> | ntc<sup>ms771</sup>/ntc<sup>ms771</sup> |
| ntc<sup>-/Df</sup> | ntc<sup>ms771</sup>/Df(3L)Exel6097 |
| ntc<sup>-/-</sup>, ntc<sup>rescue</sup> | UAS-ntc; da-GAL4, ntc<sup>ms771</sup>/ntc<sup>ms771</sup> |
| **C** |  |
| Control | w<sup>1118</sup> |
| ntc<sup>-/-</sup> | ntc<sup>ms771</sup>/ntc<sup>ms771</sup> |
| ntc<sup>-/-</sup>, ntc<sup>rescue</sup> | UAS-ntc; da-GAL4, ntc<sup>ms771</sup>/ntc<sup>ms771</sup> |
| **D** |  |
| Control | w<sup>1118</sup> |
| ntc<sup>-/-</sup> | ntc<sup>ms771</sup>/ntc<sup>ms771</sup> |
| **E** |  |
| Control | UAS-LacZ/+; da-GAL4/+ |
| ntc<sup>-/-</sup> | ntc<sup>ms771</sup>/ntc<sup>ms771</sup> |
| ntc<sup>-/Df</sup> | ntc<sup>ms771</sup>/Df(3L)Exel6097 |
| **F** |  |
| Control | w<sup>1118</sup> |
| ntc<sup>-/-</sup> | ntc<sup>ms771</sup>/ntc<sup>ms771</sup> |
| **G** |  |
| Control | w<sup>1118</sup> |
| ntc<sup>-/-</sup> | ntc<sup>ms771</sup>/ntc<sup>ms771</sup> |
| **H** |  |
| Control | w<sup>1118</sup> |
| ntc<sup>-/-</sup> | ntc<sup>ms771</sup>/ntc<sup>ms771</sup> |
| ntc<sup>-/-</sup>, ntc<sup>rescue</sup> | UAS-ntc; da-GAL4, ntc<sup>ms771</sup>/ntc<sup>ms771</sup> |
| **I** |  |
| Control | w<sup>1118</sup> |
| ntc<sup>-/-</sup> | ntc<sup>ms771</sup>/ntc<sup>ms771</sup> |

| Figure 3 |  |
| --- | --- |
| Label | Genotype |
| **A** |  |
| Control | mito-QC/UAS-LacZ<sup>RNAi</sup>; nSyb-GAL4/+ |
| ntc<sup>-/-</sup> | mito-QC/UAS-LacZ<sup>RNAi</sup>; nSyb-GAL4, ntc<sup>ms771</sup>/ntc<sup>ms771</sup> |
| ntc<sup>-/-</sup>, ntc<sup>rescue</sup> | mito-QC/UAS-ntc; nSyb-GAL4, ntc<sup>ms771</sup>/ntc<sup>ms771</sup> |
| **B** |  |
| Control | mito-QC/UAS-LacZ<sup>RNAi</sup>; nSyb-GAL4/+ |
| ntc<sup>-/-</sup> | mito-QC/UAS-LacZ<sup>RNAi</sup>; nSyb-GAL4, ntc<sup>ms771</sup>/ntc<sup>ms771</sup> |
| ntc<sup>-/-</sup>, ntc<sup>rescue</sup> | mito-QC/UAS-ntc; nSyb-GAL4, ntc<sup>ms771</sup>/ntc<sup>ms771</sup> |
| **C** |  |
| Control | mito-QC/UAS-LacZ<sup>RNAi</sup>; nSyb-GAL4/+ |
| ntc O/E | mito-QC/UAS-ntc; nSyb-GAL4/+ |
| **D** |  |
| Control | mito-QC/UAS-LacZ<sup>RNAi</sup>; nSyb-GAL4/+ |
| ntc O/E | mito-QC/UAS-ntc; nSyb-GAL4/+ |
| **E** |  |
| Control | mito-QC/UAS-LacZ<sup>RNAi</sup>; nSyb-GAL4/+ |
| USP30<sup>RNAi</sup> | mito-QC/UAS-USP30<sup>RNAi</sup>; nSyb-GAL4/+ |
| USP30<sup>RNAi</sup>, ntc<sup>-/-</sup> | mito-QC/UAS-USP30<sup>RNAi</sup>; nSyb-GAL4, ntc<sup>ms771</sup>/ntc<sup>ms771</sup> |
| **F** |  |
| Control | mito-QC/UAS-LacZ<sup>RNAi</sup>; nSyb-GAL4/+ |
| USP30<sup>RNAi</sup> | mito-QC/UAS-USP30<sup>RNAi</sup>; nSyb-GAL4/+ |
| USP30<sup>RNAi</sup>, ntc<sup>-/-</sup> | mito-QC/UAS-USP30<sup>RNAi</sup>; nSyb-GAL4, ntc<sup>ms771</sup>/ntc<sup>ms771</sup> |
| **G** |  |
| Control | Act-GAL4/UAS-LacZ |
| USP30 O/E + LacZ | Act-GAL4, UAS-USP30/UAS-LacZ |
| USP30 O/E + ntc O/E | Act-GAL4, UAS-USP30/UAS-ntc |

| Figure 4 |  |
| --- | --- |
| <u>Label</u> | <u>Genotype</u> |
| **A** |  |
| Control | mito-QC/UAS-LacZ<sup>RNAi</sup>; nSyb-GAL4/+ |
| park<sup>-/-</sup> | mito-QC/UAS-LacZ<sup>RNAi</sup>; nSyb-GAL4, park<sup>25</sup>/park<sup>25</sup> |
| park<sup>-/-</sup>, ntc O/E | mito-QC/UAS-ntc; nSyb-GAL4, park<sup>25</sup>/park<sup>25</sup> |
| **B** |  |
| Control | mito-QC/UAS-LacZ<sup>RNAi</sup>; nSyb-GAL4/+ |
| park<sup>-/-</sup> | mito-QC/UAS-LacZ<sup>RNAi</sup>; nSyb-GAL4, park<sup>25</sup>/park<sup>25</sup> |
| park<sup>-/-</sup>, ntc O/E | mito-QC/UAS-ntc; nSyb-GAL4, park<sup>25</sup>/park<sup>25</sup> |
| **C** |  |
| Control | mito-QC/UAS-LacZ<sup>RNAi</sup>; nSyb-GAL4/+ |
| Pink1<sup>-</sup> | Pink1<sup>B9</sup>/Y; mito-QC/UAS-LacZ<sup>RNAi</sup>; nSyb-GAL4/+ |
| Pink1<sup>-</sup>, ntc O/E | Pink1<sup>B9</sup>/Y; mito-QC/UAS-ntc; nSyb-GAL4/+ |
| **D** |  |
| Control | mito-QC/UAS-LacZ<sup>RNAi</sup>; nSyb-GAL4/+ |
| Pink1<sup>-</sup> | Pink1<sup>B9</sup>/Y; mito-QC/UAS-LacZ<sup>RNAi</sup>; nSyb-GAL4/+ |
| Pink1<sup>-</sup>, ntc O/E | Pink1<sup>B9</sup>/Y; mito-QC/UAS-ntc; nSyb-GAL4/+ |

| Figure 5 |  |
| --- | --- |
| <u>Label</u> | <u>Genotype</u> |
| **A** |  |
| Control | mito-QC/UAS-LacZ<sup>RNAi</sup>; nSyb-GAL4/+ |
| park<sup>-/-</sup> | mito-QC/UAS-LacZ<sup>RNAi</sup>; nSyb-GAL4, park<sup>25</sup>/park<sup>25</sup> |
| park<sup>-/-</sup>, USP30<sup>RNAi</sup> | mito-QC/UAS-USP30<sup>RNAi</sup>; nSyb-GAL4, park<sup>25</sup>/park<sup>25</sup> |
| park<sup>-/-</sup>, USP30<sup>RNAi</sup>, ntc<sup>-/-</sup> | mito-QC/UAS-USP30<sup>RNAi</sup>; nSyb-GAL4, park<sup>25</sup>,<br>PBac{WH}CG10855<sup>f07259</sup>/ntc<sup>ms771</sup> |
| **B** |  |
| Control | mito-QC/UAS-LacZ<sup>RNAi</sup>; nSyb-GAL4/+ |
| park<sup>-/-</sup> | mito-QC/UAS-LacZ<sup>RNAi</sup>; nSyb-GAL4, park<sup>25</sup>/park<sup>25</sup> |
| park<sup>-/-</sup>, USP30<sup>RNAi</sup> | mito-QC/UAS-USP30<sup>RNAi</sup>; nSyb-GAL4, park<sup>25</sup>/park<sup>25</sup> |
| park<sup>-/-</sup>, USP30<sup>RNAi</sup>, ntc<sup>-/-</sup> | mito-QC/UAS-USP30<sup>RNAi</sup>; nSyb-GAL4, park<sup>25</sup>,<br>PBac{WH}CG10855<sup>f07259</sup>/ntc<sup>ms771</sup> |
| **C** |  |
| Control | mito-QC/ UAS-LacZ<sup>RNAi</sup>; nSyb-GAL4/+ |
| Pink1<sup>-</sup> | Pink1<sup>B9</sup>/Y; mito-QC/UAS-LacZ<sup>RNAi</sup>; nSyb-GAL4/+ |
| Pink1<sup>-</sup>, USP30<sup>RNAi</sup> | Pink1<sup>B9</sup>/Y; mito-QC/UAS-USP30<sup>RNAi</sup>; nSyb-GAL4/+ |
| Pink1<sup>-</sup>, USP30<sup>RNAi</sup>, ntc<sup>-/-</sup> | Pink1<sup>B9</sup>/Y; mito-QC/UAS-USP30<sup>RNAi</sup>; nSyb-GAL4,<br>ntc<sup>ms771</sup>/ntc<sup>ms771</sup> |
| **D** |  |
| Control | mito-QC/ UAS-LacZ<sup>RNAi</sup>; nSyb-GAL4/+ |
| Pink1<sup>-</sup> | Pink1<sup>B9</sup>/Y; mito-QC/UAS-LacZ<sup>RNAi</sup>; nSyb-GAL4/+ |
| Pink1<sup>-</sup>, USP30<sup>RNAi</sup> | Pink1<sup>B9</sup>/Y; mito-QC/UAS-USP30<sup>RNAi</sup>; nSyb-GAL4/+ |
| Pink1<sup>-</sup>, USP30<sup>RNAi</sup>, ntc<sup>-/-</sup> | Pink1<sup>B9</sup>/Y; mito-QC/UAS-USP30<sup>RNAi</sup>; nSyb-GAL4,<br>ntc<sup>ms771</sup>/ntc<sup>ms771</sup> |

| **Figure 6** |  |
| --- | --- |
| <u>Label</u> | <u>Genotype</u> |
| **A** |  |
| Control | UAS-LacZ/+; da-GAL4/+ |
| USP30<sup>RNAi</sup> | UAS-USP30<sup>RNAi</sup>/+; da-GAL4/+ |
| ntc O/E | UAS-ntc/+; da-GAL4/+ |
| ntc<sup>-/-</sup> | ntc<sup>ms771</sup>/ntc<sup>ms771</sup> |
| **B** |  |
| Control | UAS-LacZ/+; da-GAL4/+ |
| USP30<sup>RNAi</sup> | UAS-USP30<sup>RNAi</sup>/+; da-GAL4/+ |
| ntc O/E | UAS-ntc/+; da-GAL4/+ |
| ntc<sup>-/-</sup> | ntc<sup>ms771</sup>/ntc<sup>ms771</sup> |
| **C** |  |
| Control | UAS-LacZ/+; da-GAL4/+ |
| USP30<sup>RNAi</sup> | UAS-USP30<sup>RNAi</sup>/+; da-GAL4/+ |
| ntc O/E | UAS-ntc/+; da-GAL4/+ |
| ntc<sup>-/-</sup> | ntc<sup>ms771</sup>/ntc<sup>ms771</sup> |
| **D** |  |
| Control | UAS-LacZ/+; da-GAL4/+ |
| USP30<sup>RNAi</sup> | UAS-USP30<sup>RNAi</sup>/+; da-GAL4/+ |
| ntc O/E | UAS-ntc/+; da-GAL4/+ |
| ntc<sup>-/-</sup> | ntc<sup>ms771</sup>/ntc<sup>ms771</sup> |
| **E** |  |
| Control | UAS-GFP-mCherry-Atg8a/UAS-LacZ<sup>RNAi</sup>/+; nSyb-GAL4/+ |
| USP30<sup>RNAi</sup> | UAS-GFP-mCherry-Atg8a/UAS-USP30<sup>RNAi</sup>/+; nSyb-GAL4/+ |
| ntc O/E | UAS-GFP-mCherry-Atg8a/UAS-ntc/+; nSyb-GAL4/+ |
| ntc<sup>-/-</sup> | UAS-GFP-mCherry-Atg8a/UAS-LacZ<sup>RNAi</sup>/+; nSyb-GAL4,<br>ntc<sup>ms771</sup>/ntc<sup>ms771</sup> |
| **F** |  |
| Control | UAS-GFP-mCherry-Atg8a/UAS-LacZ<sup>RNAi</sup>/+; nSyb-GAL4/+ |
| USP30<sup>RNAi</sup> | UAS-GFP-mCherry-Atg8a/UAS-USP30<sup>RNAi</sup>/+; nSyb-GAL4/+ |
| ntc O/E | UAS-GFP-mCherry-Atg8a/UAS-ntc/+; nSyb-GAL4/+ |
| ntc<sup>-/-</sup> | UAS-GFP-mCherry-Atg8a/UAS-LacZ<sup>RNAi</sup>/+; nSyb-GAL4,<br>ntc<sup>ms771</sup>/ntc<sup>ms771</sup> |

| Figure 7 |  |
| --- | --- |
| <u>Label</u> | <u>Genotype</u> |
| **A** |  |
| Control | UAS-LacZ/+; da-GAL4/+ |
| ntc O/E | UAS-ntc/+; da-GAL4/+ |
| USP30<sup>RNAi</sup> | UAS-USP30<sup>RNAi</sup>/+; da-GAL4/+ |
| **B** |  |
| Control | UAS-LacZ/+; da-GAL4/+ |
| ntc O/E | UAS-ntc/+; da-GAL4/+ |
| USP30<sup>RNAi</sup> | UAS-USP30<sup>RNAi</sup>/+; da-GAL4/+ |
| **C** |  |
| Control | w<sup>1118</sup> |
| park<sup>-/-</sup> | park<sup>25</sup>/park<sup>25</sup> |
| ntc<sup>-/-</sup> | ntc<sup>ms771</sup>/PBac{WH}CG10855<sup>f07259</sup> |
| park<sup>-/-</sup>, ntc<sup>-/-</sup> | park<sup>25</sup>, ntc<sup>ms771</sup>/park<sup>25</sup>, PBac{WH}CG10855<sup>f07259</sup> |
| **D** |  |
| Control | w<sup>1118</sup> |
| park<sup>-/-</sup> | park<sup>25</sup>/park<sup>25</sup> |
| ntc<sup>-/-</sup> | ntc<sup>ms771</sup>/PBac{WH}CG10855<sup>f07259</sup> |
| park<sup>-/-</sup>, ntc<sup>-/-</sup> | park<sup>25</sup>, ntc<sup>ms771</sup>/park<sup>25</sup>, PBac{WH}CG10855<sup>f07259</sup> |
| **E** |  |
| Control | UAS-LacZ/+; da-GAL4/+ |
| park<sup>-/-</sup> | UAS-LacZ/+; da-GAL4, park<sup>25</sup>/park<sup>25</sup> |
| park<sup>-/-</sup>; ntc O/E | UAS-ntc/+; da-GAL4, park<sup>25</sup>/park<sup>25</sup> |
| **F** |  |
| Control | UAS-LacZ/+; da-GAL4/+ |
| park<sup>-/-</sup> | UAS-LacZ/+; da-GAL4, park<sup>25</sup>/park<sup>25</sup> |
| park<sup>-/-</sup>; ntc O/E | UAS-ntc/+; da-GAL4, park<sup>25</sup>/park<sup>25</sup> |

| Supplementary Figure 1 |  |
| --- | --- |
| <u>Label</u> | <u>Genotype</u> |
| **A** |  |
| Control | mtx-QC/UAS-LacZ<sup>RNAi</sup>; nSyb-GAL4/+ |
| USP30<sup>RNAi</sup> | mtx-QC/UAS-USP30<sup>RNAi</sup>; nSyb-GAL4/+ |
| ntc | mtx-QC/UAS-ntc; nSyb-GAL4/+ |
| ntc<sup>-/-</sup> | mtx-QC/UAS-LacZ<sup>RNAi</sup>; nSyb-GAL4, ntc<sup>ms771</sup>/ntc<sup>ms771</sup> |
| **B** |  |
| Control | mtx-QC/UAS-LacZ<sup>RNAi</sup>; nSyb-GAL4/+ |
| USP30<sup>RNAi</sup> | mtx-QC/UAS-USP30<sup>RNAi</sup>; nSyb-GAL4/+ |
| ntc | mtx-QC/UAS-ntc; nSyb-GAL4/+ |
| ntc<sup>-/-</sup> | mtx-QC/UAS-LacZ<sup>RNAi</sup>; nSyb-GAL4, ntc<sup>ms771</sup>/ntc<sup>ms771</sup> |

| Supplementary Figure 2 |  |
| --- | --- |
| <u>Label</u> | <u>Genotype</u> |
| **A** |  |
| Control | mito-QC/UAS-LacZ<sup>RNAi</sup>; nSyb-GAL4/+ |
| FBXO7 O/E | mito-QC/UAS-FBXO7; nSyb-GAL4/+ |
| **B** |  |
| Control | mito-QC/UAS-LacZ<sup>RNAi</sup>; nSyb-GAL4/+ |
| FBXO7 O/E | mito-QC/UAS-FBXO7; nSyb-GAL4/+ |

| Supplementary Figure 3 |  |
| --- | --- |
| <u>Label</u> | <u>Genotype</u> |
| **A** |  |
| Control | Mef2-GAL4/UAS-LacZ<sup>RNAi</sup>; mito-QC/+ |
| USP30<sup>RNAi</sup> | Mef2-GAL4/UAS-USP30<sup>RNAi</sup>; mito-QC/+ |
| **B** |  |
| Control | Mef2-GAL4/UAS-LacZ<sup>RNAi</sup>; mito-QC/+ |
| USP30<sup>RNAi</sup> | Mef2-GAL4/UAS-USP30<sup>RNAi</sup>; mito-QC/+ |
| **C** |  |
| Control | UAS-LacZ<sup>RNAi</sup>/+; da-GAL4/+ |
| USP30<sup>RNAi</sup> | UAS-USP30<sup>RNAi</sup>/+; da-GAL4/+ |
| Control | UAS-LacZ<sup>RNAi</sup>/+; nSyb-GAL4/+ |
| USP30<sup>RNAi</sup> | UAS-USP30<sup>RNAi</sup>/+; nSyb-GAL4/+ |
| **D** |  |
| Control | Act-GAL4/UAS-LacZ |
| USP30 O/E + LacZ | Act-GAL4, UAS-USP30/UAS-LacZ |
| USP30 O/E + parkin O/E | Act-GAL4, UAS-USP30/UAS-parkin |
| USP30 O/E + March5 O/E | Act-GAL4, UAS-USP30/+; UAS-March5/+ |
| USP30 O/E + Mul1 O/E | Act-GAL4, UAS-USP30/+; UAS-Mul1/+ |
| **E** |  |
| Control | mito-QC/UAS-LacZ<sup>RNAi</sup>; nSyb-GAL4/+ |
| parkin O/E | mito-QC/UAS- parkin<sup>c2</sup>; nSyb-GAL4/+ |
| **F** |  |
| Control | mito-QC/UAS-LacZ<sup>RNAi</sup>; nSyb-GAL4/+ |
| parkin O/E | mito-QC/UAS- parkin<sub>C2</sub>; nSyb-GAL4/+ |
| **G** |  |
| Control | mito-QC/+; nSyb-GAL4/+ |
| Mul1<sup>-/-</sup> | mito-QC/+; nSyb-GAL4, Mul1[A6]/Mul1[A6] |
| **H** |  |
| Control | mito-QC/+; nSyb-GAL4/+ |
| Mul1<sup>-/-</sup> | mito-QC/+; nSyb-GAL4, Mul1[A6]/Mul1[A6] |
| **I** |  |
| Control | mito-QC/UAS-LacZ<sup>RNAi</sup>; nSyb-GAL4/+ |
| March5<sup>RNAi KK</sup> | mito-QC/UAS-March5<sup>RNAi KK</sup>; nSyb-GAL4/+ |
| March5<sup>RNAi GD</sup> | mito-QC/UAS-March5<sup>RNAi GD</sup>; nSyb-GAL4/+ |
| **J** |  |
| Control | mito-QC/UAS-LacZ<sup>RNAi</sup>; nSyb-GAL4/+ |
| March5<sup>RNAi KK</sup> | mito-QC/UAS-March5<sup>RNAi KK</sup>; nSyb-GAL4/+ |
| March5<sup>RNAi GD</sup> | mito-QC/UAS-March5<sup>RNAi GD</sup>; nSyb-GAL4/+ |

| Supplementary Figure 4 |  |
| --- | --- |
| <u>Label</u> | <u>Genotype</u> |
| **A** |  |
| Control | mito-QC/UAS-LacZ<sup>RNAi</sup>; nSyb-GAL4/+ |
| Atg8a<sup>-/-</sup>, USP30<sup>RNAi</sup> | Atg8a[KG07569]/Y; mito-QC/UAS-USP30<sup>RNAi</sup>; nSyb-GAL4/+ |
| Atg8a<sup>-/-</sup>, ntc O/E | Atg8a[KG07569]/Y; mito-QC/UAS-ntc; nSyb-GAL4/+ |
| Atg8a<sup>-/-</sup>, ntc<sup>-/-</sup> | Atg8a[KG07569]/Y; mito-QC/ LacZ<sup>RNAi</sup>; nSyb-GAL4, ntc<sup>ms771</sup>/ntc<sup>ms771</sup> |
| **B** |  |
| Control | mito-QC/UAS-LacZ<sup>RNAi</sup>; nSyb-GAL4/+ |
| Atg8a<sup>-/-</sup>, USP30<sup>RNAi</sup> | Atg8a[KG07569]/Y; mito-QC/UAS-USP30<sup>RNAi</sup>; nSyb-GAL4/+ |
| Atg8a<sup>-/-</sup>, ntc O/E | Atg8a[KG07569]/Y; mito-QC/UAS-ntc; nSyb-GAL4/+ |
| Atg8a<sup>-/-</sup>, ntc<sup>-/-</sup> | Atg8a[KG07569]/Y; mito-QC/ LacZ<sup>RNAi</sup>; nSyb-GAL4, ntc<sup>ms771</sup>/ntc<sup>ms771</sup> |
| **C** |  |
| Control | mito-QC/UAS-LacZ<sup>RNAi</sup>; nSyb-GAL4/+ |
| park<sup>-/-</sup> | mito-QC/UAS-LacZ<sup>RNAi</sup>; nSyb-GAL4, park<sup>25</sup>/park<sup>25</sup> |
| USP30<sup>RNAi</sup>, park<sup>-/-</sup> | mito-QC/UAS-USP30<sup>RNAi</sup>; nSyb-GAL4, park<sup>25</sup>/park<sup>25</sup> |
| Atg8a<sup>-/-</sup>, USP30<sup>RNAi</sup>, park<sup>-/-</sup> | Atg8a[KG07569]/Y; mito-QC/UAS-USP30<sup>RNAi</sup>; nSyb-GAL4, park<sup>25</sup>/park<sup>25</sup> |
| **D** |  |
| Control | mito-QC/UAS-LacZ<sup>RNAi</sup>; nSyb-GAL4/+ |
| park<sup>-/-</sup> | mito-QC/UAS-LacZ<sup>RNAi</sup>; nSyb-GAL4, park<sup>25</sup>/park<sup>25</sup> |
| USP30<sup>RNAi</sup>, park<sup>-/-</sup> | mito-QC/UAS-USP30<sup>RNAi</sup>; nSyb-GAL4, park<sup>25</sup>/park<sup>25</sup> |
| Atg8a<sup>-/-</sup>, USP30<sup>RNAi</sup>, park<sup>-/-</sup> | Atg8a[KG07569]/Y; mito-QC/UAS-USP30<sup>RNAi</sup>; nSyb-GAL4, park<sup>25</sup>/park<sup>25</sup> |

| **Supplementary Figure 5** |  |
| --- | --- |
| **<u>Label</u>** | **<u>Genotype</u>** |
| **A** |  |
| Control | UAS-LacZ/+; da-GAL4/+ |
| ntc O/E | UAS-ntc/+; da-GAL4/+ |
| USP30<sup>RNAi</sup> | UAS-USP30<sup>RNAi</sup>/+; da-GAL4/+ |
| **B** |  |
| Control | UAS-LacZ/+; da-GAL4/+ |
| ntc O/E | UAS-ntc/+; da-GAL4/+ |
| USP30<sup>RNAi</sup> | UAS-USP30<sup>RNAi</sup>/+; da-GAL4/+ |

| C |  |
| --- | --- |
| Control | w<sup>1118</sup> |
| ntc<sup>-/-</sup> | ntc<sup>ms771</sup>/ntc<sup>ms771</sup> |

## References

1. Yamano K, Youle RJ. PINK1 is degraded through the N-end rule pathway. Autophagy. 2013;9(11):1758–69.

2. Okatsu K, Kimura M, Oka T, Tanaka K, Matsuda N. Unconventional PINK1 localization to the outer membrane of depolarized mitochondria drives Parkin recruitment. J Cell Sci. 2015;128(5):964–78.

3. Okatsu K, Koyano F, Kimura M, Kosako H, Saeki Y, Tanaka K, et al. Phosphorylated ubiquitin chain is the genuine Parkin receptor. J Cell Biol. 2015;209(1):111–28.

4. Gladkova C, Maslen SL, Skehel JM, Komander D. Mechanism of parkin activation by PINK1. Nature. 2018;559(7714):410–4.

5. Sauve V, Sung G, Soya N, Kozlov G, Blaimschein N, Miotto LS, et al. Mechanism of parkin activation by phosphorylation. Nature structural & molecular biology. 2018;25(7):623–30.

6. Ordureau A, Sarraf SA, Duda DM, Heo JM, Jedrychowski MP, Sviderskiy VO, et al. Quantitative proteomics reveal a feedforward mechanism for mitochondrial PARKIN translocation and ubiquitin chain synthesis. Mol Cell. 2014;56(3):360–75.

7. Bingol B, Sheng M. Mechanisms of mitophagy: PINK1, Parkin, USP30 and beyond. Free Radic Biol Med. 2016;100:210–22.

8. Lazarou M, Sliter DA, Kane LA, Sarraf SA, Wang C, Burman JL, et al. The ubiquitin kinase PINK1 recruits autophagy receptors to induce mitophagy. Nature. 2015;524(7565):309–14.

9. Yamano K, Kikuchi R, Kojima W, Hayashida R, Koyano F, Kawawaki J, et al. Critical role of mitochondrial ubiquitination and the OPTN-ATG9A axis in mitophagy. J Cell Biol. 2020;219(9).

10. Pickrell AM, Youle RJ. The roles of PINK1, parkin, and mitochondrial fidelity in Parkinson’s disease. Neuron. 2015;85(2):257–73.

11. Clark IE, Dodson MW, Jiang C, Cao JH, Huh JR, Seol JH, et al. Drosophila pink1 is required for mitochondrial function and interacts genetically with parkin. Nature. 2006;441(7097):1162–6.

12. Park J, Lee SB, Lee S, Kim Y, Song S, Kim S, et al. Mitochondrial dysfunction in Drosophila PINK1 mutants is complemented by parkin. Nature. 2006;441(7097):1157–61.

13. Greene JC, Whitworth AJ, Kuo I, Andrews LA, Feany MB, Pallanck LJ. Mitochondrial pathology and apoptotic muscle degeneration in Drosophila parkin mutants. Proceedings of the National Academy of Sciences of the United States of America. 2003;100(7):4078–83.

14. Whitworth AJ, Theodore DA, Greene JC, Benes H, Wes PD, Pallanck LJ. Increased glutathione S-transferase activity rescues dopaminergic neuron loss in a Drosophila model of Parkinson’s disease. Proc Natl Acad Sci U S A. 2005;102(22):8024–9.

15. Poole AC, Thomas RE, Andrews LA, McBride HM, Whitworth AJ, Pallanck LJ. The PINK1/Parkin pathway regulates mitochondrial morphology. Proceedings of the National Academy of Sciences of the United States of America. 2008;105(5):1638–43.

16. Ziviani E, Tao RN, Whitworth AJ. Drosophila parkin requires PINK1 for mitochondrial translocation and ubiquitinates mitofusin. Proceedings of the National Academy of Sciences of the United States of America. 2010;107(11):5018–23.

17. Vincow ES, Merrihew G, Thomas RE, Shulman NJ, Beyer RP, MacCoss MJ, et al. The PINK1-Parkin pathway promotes both mitophagy and selective respiratory chain turnover in vivo. Proceedings of the National Academy of Sciences of the United States of America. 2013;110(16):6400–5.

18. Cornelissen T, Verstreken P, Vandenberghe W. Imaging mitophagy in the fruit fly. Autophagy. 2018;14(9):1656–7.

19. Kim YY, Um JH, Yoon JH, Kim H, Lee DY, Lee YJ, et al. Assessment of mitophagy in mt-Keima Drosophila revealed an essential role of the PINK1-Parkin pathway in mitophagy induction in vivo. FASEB J. 2019;33(9):9742–51.

20. Lee JJ, Sanchez-Martinez A, Martinez Zarate A, Beninca C, Mayor U, Clague MJ, et al. Basal mitophagy is widespread in Drosophila but minimally affected by loss of Pink1 or parkin. J Cell Biol. 2018;217(5):1613–22.

21. McWilliams TG, Prescott AR, Montava-Garriga L, Ball G, Singh F, Barini E, et al. Basal Mitophagy Occurs Independently of PINK1 in Mouse Tissues of High Metabolic Demand. Cell metabolism. 2018;27(2):439–49 e5.

22. Di Fonzo A, Dekker MC, Montagna P, Baruzzi A, Yonova EH, Correia Guedes L, et al. FBXO7 mutations cause autosomal recessive, early-onset parkinsonian-pyramidal syndrome. Neurology. 2009;72(3):240–5.

23. Zhou ZD, Lee JCT, Tan EK. Pathophysiological mechanisms linking F-box only protein 7 (FBXO7) and Parkinson’s disease (PD). Mutat Res Rev Mutat Res. 2018;778:72–8.

24. Skowyra D, Craig KL, Tyers M, Elledge SJ, Harper JW. F-box proteins are receptors that recruit phosphorylated substrates to the SCF ubiquitin-ligase complex. Cell. 1997;91(2):209–19.

25. Hsu JM, Lee YC, Yu CT, Huang CY. Fbx7 functions in the SCF complex regulating Cdk1-cyclin B-phosphorylated hepatoma up-regulated protein (HURP) proteolysis by a proline-rich region. J Biol Chem. 2004;279(31):32592–602.

26. Kirk R, Laman H, Knowles PP, Murray-Rust J, Lomonosov M, Meziane el K, et al. Structure of a conserved dimerization domain within the F-box protein Fbxo7 and the PI31 proteasome inhibitor. The Journal of biological chemistry. 2008;283(32):22325–35.

27. Burchell VS, Nelson DE, Sanchez-Martinez A, Delgado-Camprubi M, Ivatt RM, Pogson JH, et al. The Parkinson’s disease-linked proteins Fbxo7 and Parkin interact to mediate mitophagy. Nat Neurosci. 2013;16(9):1257–65.

28. Bader M, Arama E, Steller H. A novel F-box protein is required for caspase activation during cellular remodeling in Drosophila. Development. 2010;137(10):1679–88.

29. Bader M, Benjamin S, Wapinski OL, Smith DM, Goldberg AL, Steller H. A conserved F box regulatory complex controls proteasome activity in Drosophila. Cell. 2011;145(3):371–82.

30. Song S, Jang S, Park J, Bang S, Choi S, Kwon KY, et al. Characterization of PINK1 (PTEN-induced putative kinase 1) mutations associated with Parkinson disease in mammalian cells and Drosophila. J Biol Chem. 2013;288(8):5660–72.

31. Whitworth AJ, Lee JR, Ho VM, Flick R, Chowdhury R, McQuibban GA. Rhomboid-7 and HtrA2/Omi act in a common pathway with the Parkinson’s disease factors Pink1 and Parkin. Dis Model Mech. 2008;1(2-3):168–74; discussion 73.

32. Riparbelli MG, Callaini G. The Drosophila parkin homologue is required for normal mitochondrial dynamics during spermiogenesis. Dev Biol. 2007;303(1):108–20.

33. Meulener M, Whitworth AJ, Armstrong-Gold CE, Rizzu P, Heutink P, Wes PD, et al. Drosophila DJ-1 mutants are selectively sensitive to environmental toxins associated with Parkinson’s disease. Curr Biol. 2005;15(17):1572–7.

34. Malik BR, Godena VK, Whitworth AJ. VPS35 pathogenic mutations confer no dominant toxicity but partial loss of function in Drosophila and genetically interact with parkin. Hum Mol Genet. 2015;24(21):6106–17.

35. Allen GF, Toth R, James J, Ganley IG. Loss of iron triggers PINK1/Parkin-independent mitophagy. EMBO Rep. 2013;14(12):1127–35.

36. Shen JL, Fortier TM, Wang R, Baehrecke EH. Vps13D functions in a Pink1-dependent and Parkin-independent mitophagy pathway. J Cell Biol. 2021;220(11).

37. Montava-Garriga L, Singh F, Ball G, Ganley IG. Semi-automated quantitation of mitophagy in cells and tissues. Mech Ageing Dev. 2020;185:111196.

38. Marcassa E, Kallinos A, Jardine J, Rusilowicz-Jones EV, Martinez A, Kuehl S, et al. Dual role of USP30 in controlling basal pexophagy and mitophagy. EMBO Rep. 2018;19(7).

39. Ordureau A, Paulo JA, Zhang J, An H, Swatek KN, Cannon JR, et al. Global Landscape and Dynamics of Parkin and USP30-Dependent Ubiquitylomes in iNeurons during Mitophagic Signaling. Mol Cell. 2020;77(5):1124–42 e10.

40. Rusilowicz-Jones EV, Jardine J, Kallinos A, Pinto-Fernandez A, Guenther F, Giurrandino M, et al. USP30 sets a trigger threshold for PINK1-PARKIN amplification of mitochondrial ubiquitylation. Life Sci Alliance. 2020;3(8).

41. Bingol B, Tea JS, Phu L, Reichelt M, Bakalarski CE, Song Q, et al. The mitochondrial deubiquitinase USP30 opposes parkin-mediated mitophagy. Nature. 2014;510(7505):370–5.

42. Phu L, Rose CM, Tea JS, Wall CE, Verschueren E, Cheung TK, et al. Dynamic Regulation of Mitochondrial Import by the Ubiquitin System. Mol Cell. 2020;77(5):1107–23 e10.

43. Yun J, Puri R, Yang H, Lizzio MA, Wu C, Sheng ZH, et al. MUL1 acts in parallel to the PINK1/parkin pathway in regulating mitofusin and compensates for loss of PINK1/parkin. Elife. 2014;3:e01958.

44. Rojansky R, Cha MY, Chan DC. Elimination of paternal mitochondria in mouse embryos occurs through autophagic degradation dependent on PARKIN and MUL1. Elife. 2016;5.

45. Zheng J, Chen X, Liu Q, Zhong G, Zhuang M. Ubiquitin ligase MARCH5 localizes to peroxisomes to regulate pexophagy. J Cell Biol. 2022;221(1).

46. Chen Z, Liu L, Cheng Q, Li Y, Wu H, Zhang W, et al. Mitochondrial E3 ligase MARCH5 regulates FUNDC1 to fine-tune hypoxic mitophagy. EMBO Rep. 2017;18(3):495–509.

47. Koyano F, Yamano K, Kosako H, Kimura Y, Kimura M, Fujiki Y, et al. Parkin-mediated ubiquitylation redistributes MITOL/March5 from mitochondria to peroxisomes. EMBO Rep. 2019;20(12):e47728.

48. Liang JR, Martinez A, Lane JD, Mayor U, Clague MJ, Urbe S. USP30 deubiquitylates mitochondrial Parkin substrates and restricts apoptotic cell death. EMBO Rep. 2015;16(5):618–27.

49. Pan W, Wang Y, Bai X, Yin Y, Dai L, Zhou H, et al. Deubiquitinating enzyme USP30 negatively regulates mitophagy and accelerates myocardial cell senescence through antagonism of Parkin. Cell Death Discov. 2021;7(1):187.

50. Lazarou M, McKenzie M, Ohtake A, Thorburn DR, Ryan MT. Analysis of the assembly profiles for mitochondrial- and nuclear-DNA-encoded subunits into complex I. Mol Cell Biol. 2007;27(12):4228–37.

51. Nguyen TN, Padman BS, Usher J, Oorschot V, Ramm G, Lazarou M. Atg8 family LC3/GABARAP proteins are crucial for autophagosome-lysosome fusion but not autophagosome formation during PINK1/Parkin mitophagy and starvation. J Cell Biol. 2016;215(6):857–74.

52. Jipa A, Vedelek V, Merenyi Z, Urmosi A, Takats S, Kovacs AL, et al. Analysis of Drosophila Atg8 proteins reveals multiple lipidation-independent roles. Autophagy. 2021;17(9):2565–75.

53. Yoshii SR, Mizushima N. Monitoring and Measuring Autophagy. Int J Mol Sci. 2017;18(9).

54. Lorincz P, Mauvezin C, Juhasz G. Exploring Autophagy in Drosophila. Cells. 2017;6(3).

55. Mauvezin C, Ayala C, Braden CR, Kim J, Neufeld TP. Assays to monitor autophagy in Drosophila. Methods. 2014;68(1):134–9.

56. Nagy P, Karpati M, Varga A, Pircs K, Venkei Z, Takats S, et al. Atg17/FIP200 localizes to perilysosomal Ref(2)P aggregates and promotes autophagy by activation of Atg1 in Drosophila. Autophagy. 2014;10(3):453–67.

57. Low P, Varga A, Pircs K, Nagy P, Szatmari Z, Sass M, et al. Impaired proteasomal degradation enhances autophagy via hypoxia signaling in Drosophila. BMC Cell Biol. 2013;14:29.

58. Gersch M, Gladkova C, Schubert AF, Michel MA, Maslen S, Komander D. Mechanism and regulation of the Lys6-selective deubiquitinase USP30. Nature structural & molecular biology. 2017;24(11):920–30.

59. Usher JL, Sanchez-Martinez A, Terriente-Felix A, Chen PL, Lee JJ, Chen CH, et al. Parkin drives pS65-Ub turnover independently of canonical autophagy in Drosophila. EMBO Rep. 2022:e202153552.

60. Zhou ZD, Xie SP, Sathiyamoorthy S, Saw WT, Sing TY, Ng SH, et al. F-box protein 7 mutations promote protein aggregation in mitochondria and inhibit mitophagy. Hum Mol Genet. 2015;24(22):6314–30.

61. Ashrafi G, Schwarz TL. The pathways of mitophagy for quality control and clearance of mitochondria. Cell Death Differ. 2013;20(1):31–42.

62. Pickles S, Vigie P, Youle RJ. Mitophagy and Quality Control Mechanisms in Mitochondrial Maintenance. Curr Biol. 2018;28(4):R170–R85.

63. Cunningham CN, Baughman JM, Phu L, Tea JS, Yu C, Coons M, et al. USP30 and parkin homeostatically regulate atypical ubiquitin chains on mitochondria. Nature cell biology. 2015;17(2):160–9.

64. Ordureau A, Heo JM, Duda DM, Paulo JA, Olszewski JL, Yanishevski D, et al. Defining roles of PARKIN and ubiquitin phosphorylation by PINK1 in mitochondrial quality control using a ubiquitin replacement strategy. Proceedings of the National Academy of Sciences of the United States of America. 2015;112(21):6637–42.

65. Teixeira FR, Randle SJ, Patel SP, Mevissen TE, Zenkeviciute G, Koide T, et al. Gsk3beta and Tomm20 are substrates of the SCFFbxo7/PARK15 ubiquitin ligase associated with Parkinson’s disease. Biochem J. 2016;473(20):3563–80.

66. Sullivan WA, M.; Hawley, R. S. Drosophila protocols. Cold Spring Harbor Laboratory Press. 2000.

67. Pogson JH, Ivatt RM, Sanchez-Martinez A, Tufi R, Wilson E, Mortiboys H, et al. The Complex I Subunit NDUFA10 Selectively Rescues Drosophila pink1 Mutants through a Mechanism Independent of Mitophagy. PLoS genetics. 2014;10(11):e1004815.

